# Chemoproteomics Maps Glycolytic Targetome in Cancer Cells

**DOI:** 10.1101/2020.11.18.387670

**Authors:** Yang Tian, Ning Wan, Ming Ding, Chang Shao, Nian Wang, Qiuyu Bao, Wenjie Lu, Haiyang Hu, Huiyong Sun, Chenxi Yang, Kun Zhou, Shuai Chen, Guangji Wang, Hui Ye, Haiping Hao

## Abstract

Hyperactivated glycolysis, favoring uncontrolled growth and metastasis by producing essential metabolic intermediates engaging bioenergetics and biosynthesis, is a metabolic hallmark of most cancer cells. Although sporadic information has revealed glycolytic metabolites also possess non-metabolic function as signaling molecules, it remains largely elusive how these metabolites interact with and functionally regulate their binding targets. Here we introduce a Target Responsive Accessibility Profiling (TRAP) approach that measures ligand binding-induced steric hindrance in protein targets via global profiling accessibility changes in reactive lysines, and mapped 913 target candidates and 2,487 interactions for 10 major glycolytic metabolites in cancer cells via TRAP. The elucidated targetome uncovers diverse regulatory modalities of glycolytic metabolites involving the direct perturbation of carbohydrate metabolism enzymes, intervention of transcriptional control, modulation of proteome-level acetylation and protein complex assemblies. The advantages gained from glycolysis by cancer cells are expanded by discovering lactate as a ligand for an orphan transcriptional regulator TRIM 28 that promotes p53 degradation, and by identifying pyruvate acting against a cell apoptosis inducer trichostatin A via attenuating protein acetylation. Lastly, the inhibition of glycolytic key enzymes led to identify an intrinsically active glycolytic intermediate glyceraldehyde 3-phosphate that elicits its cytotoxicity by engaging with ENO1 and MTHFD1. Collectively, the glycolytic targetome depicted by TRAP constitutes a fertile resource for understanding how glycolysis finely tunes metabolism and signaling in support of cancer cells, and fostering the exploitation of glycolytic targetome as promising nodes for anti-cancer therapeutics development.

---

**E**merging evidence supports the concept that metabolic dysregulation is not only the readouts of diseases but more likely the pathological causes and driving forces of disease progression^1-3^. In agreement, endogenous metabolites are endowed with newly discovered roles including immune-mediator and epigenetic regulator other than their conventional function as metabolic fuels^4-7^. Nevertheless, we are still in the exploratory phase of gathering sporadic information about the functions and particularly the targets of bioactive metabolites to which they elicit diverse modulatory effects for finely tuning cellular signal transduction and metabolism^8-13^.

Glycolysis is a well-established hallmark of cancers and known to confer significant advantages to support cancer cell survival, growth, and invasion over normal populations. The metabolic intermediates produced in glycolysis have been substantiated to serve as building blocks for lipogenesis and macromolecule synthesis that are essential for cancer cell growth and survival^14^. Although increasing evidences indicate that glycolytic metabolites can also direct communication intertwining cancer metabolism and other signaling pathways^11,15^, the knowledge of what proteins interact with glycolytic metabolites and, therefore, how they are functionally regulated by glycolysis remains largely elusive. We thus sought to depict the landscape of glycolytic metabolites-interacting proteins, termed herein as glycolytic targetome, because of the promiscuity of endogenous metabolites in protein binding in cancer cells. The decoded glycolytic targetome network is important for illuminating new mechanisms to understand why glycolysis is reprogrammed by cancer cells to hyperactive status besides fueling ATP production and biosynthesis, and for translating such knowledge to exploitable protein targets for anti-cancer therapy.

## RESULTS

### Benchmarking the Target Responsive Accessibility Profiling Approach for Targetome Mining from Cell Milieu

Notably, a **T**arget **R**esponsive **A**ccessibility **P**rofiling (**TRAP)** approach that probes proteome-level accessibility alterations due to ligand engagement was developed for mining the targetome of glycolytic metabolites in this study (**Fig. 1a**). TRAP differs from previous target discovery approaches that measure ligand-induced stability changes of target proteins in response to proteolytic, thermal, solvent, reagent and other denaturation stresses^8,16-20^, which assume that the transient and low-affinity metabolite interactions are competent to significantly impact the stability of the binding targetome. TRAP is established on a premise that, due to ligand binding and the concomitantly increased steric hindrance, reactive residues in the ligand-bound region of target proteins are labeled by covalent chemical probes to different extents (**Fig. 1a**). Therefore, by quantifying peptides with and without ligand incubation, we can define a TRAP ratio **R**_ligand/control_ that reflects the altered accessibility of each examined reactive lysine and assign those displaying significant intensity changes as target-responsive peptides (**TRP**s) of ligands, which determine the ligand-bound target proteins and concomitantly indicate the proximal region where the engagement occurs. We first benchmarked TRAP using ribonuclease A (RNase, **Fig. 1b**) and its two ligands of different affinities, cytidine diphosphate (CDP) and cytidine triphosphate (CTP)^21^, as revealed by native mass spectrometry (**Fig. 1c**) and microscale thermophoresis (**Extended Data Fig. 1a**). A reductive dimethylation reaction mix was employed as the covalent probe that detects accessibility for free lysine ε-amine groups on RNase. Indeed, the resultant time-resolved intact mass measurement confirms retarded reaction kinetic upon ligand engagement (**Fig. 1d**). In agreement, subsequent proteolysis and quantitative analysis of these proteins pinpointed TRPs that are located in proximity to the CDP/CTP-binding pocket^21^ display significantly reduced accessibility following ligand incubation, whereas peptides that are located remotely from the binding site hold constant R_ligand/control_ (**Fig. 1e, Extended Data Fig. 1b**). Moreover, accessibility of TRPs for CTP in RNase A decreased more markedly than those for CDP, in line with their different affinities. Besides revealing the ligand-binding sites, we asked whether TRAP can also detect accessibility changes that reflect global conformational alterations elicited by allosteric regulation. PKM2 is a multi-domain metabolic enzyme responsible for the conversion of phosphoenolpyruvate (PEP) to pyruvate, whereas FBP can allosterically trigger its tetramerization (**Extended Data Fig. 1c**). Upon FBP incubation, TRAP identifies not only the exact FBP-binding site K^433^ in PKM2 that shows the greatest decrease of ion intensity (**R_FBP/control_**=0.07), but also TRPs that carry K^270^ and K^337^ (PDB 4YJ5, **Fig. 1f-g, Extended Data Fig. 1d**). Crystal structure shows the proximity of K^270^ and K^337^ to the active site, agreeing with a partially closed conformation of the active site induced by FBP activation and explains the observed accessibility decrease^22^. In contrast, K^422^ is located near the C-C’ subunit interface^22^ and is distant from the FBP binding site and the active site according to the measured Euclidean distance (**Extended Data Fig. 1c**). Its decreased accessibility thus suggests the influenced inter-subunit contacts along the C-domain by FBP-induced PKM2 conformational change. Collectively, we highlight TRAP as a complementary approach to classic biophysical crystallography in revealing native metabolite-protein interactions.

**Figure 1.**
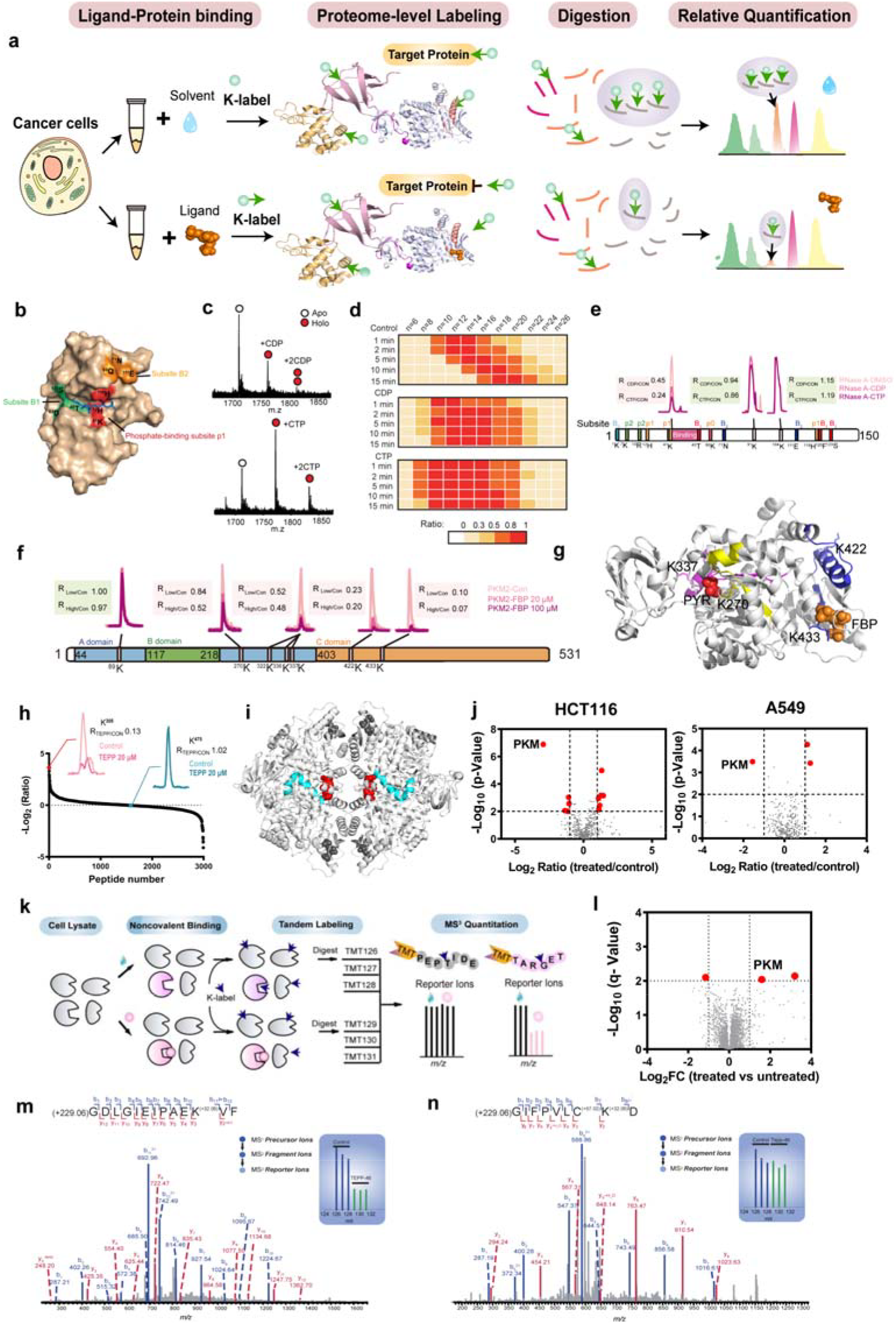
Benchmarking the TRAP approach for targetome mining in native cell milieu. **(a)** Illustration of the TRAP workflow. **(b)** Docking analysis of ribonuclease A (RNase A) with its ligand 5’-cytidine diphosphate (CDP). **(c)** Native mass spectrometry showed CDP and (5’-cytidine triphosphate) CTP bind to RNase A at different affinities. **(d)** Protein level-TRAP showed impeded mass shift of chemically labeled-RNase A in response to ligand binding. **(e)** Extracted ion chromatograms (EICs) of peptides carrying labeled lysines identify the ligand binding sites in RNase A. **(f)** TRAP identified labeled lysines that showed decreased accessibility of the active sites and the binding sites of FBP in PKM2 based on the representative EICs. **(g)** Crystallographic structure of PKM2 bound to FBP (orange) and pyruvate (red) (PDB: 4YJ5). The TRPs of FBP containing K^270^ (yellow), K337 (magenta) and K^422^/K^433^ (blue) are color-coded, respectively. **(h)** TRAP Ratio_TEPP-46/Control_ and EICs (inserts) of peptides carrying labeled lysines identify the binding sites of TEPP-46 in PKM2. **(i)** X-ray structure validates TRAP-identified binding sites of TEPP-46 (red, sphere) in PKM2 (PDB: 3U2Z). The identified TRP covering 295-316 are colored in blue. **(j)** Volcano plots identify the binding target PKM2 for TEPP-46 (10 μM) from HCT116 and A549 cell lysates (n=5). Proteins displaying significantly altered accessibility change following TEPP-46 administration with the significance cutoff of p<0.01 and the TRAP ratio (treated/control) cutoff of >2 or <0.5 are highlighted in red. **(k)** Multiplexed-TRAP workflow is empowered by 6-plex TMT isobaric labeling reagents. **(l)** Volcano plot of the multiplexed-TRAP experiments for TEPP-46 (10 μM) from the HCT116 cell lysates (n=3). Proteins displaying significantly altered accessibility change following TEPP-46 administration with the significance cutoff of q<0.01 and the TRAP ratio (treated/control) cutoff of >2 or <0.5 are highlighted in red. **(m)** MS3-based TMT reporter ions identified the TRP from PKM2 based on the significantly decreased accessibility in response to TEPP-46 binding. **(n)** A representative non-TRP from PKM2 displayed negligible change in accessibility following TEPP-46 incubation.

Since it remains less explored whether approaches assessing protein accessibility changes upon ligand binding are amenable to cellular systems, we next set out to leverage TRAP in identifying binding targets for ligands in complex cell milieu and ultimately in mapping the glycolytic metabolite targetome. We first used TEPP-46, an anticancer reagent known to specifically bind to and activate PKM2 (AC_50_ value ~92 nM), as a model ligand^23^. We prepared lysates of HCT116 cells in nondenaturing condition, labeled the proteome, and applied label free quantification (LFQ) proteomics workflow to determine the accessibility changes of quantified lysines via the TRAP workflow. Remarkably, the TRP that displays the most pronouncedly changed accessibility harbors K^305^ of PKM2 (**Fig. 1h**). This indicates the tightest engagement of TEPP-46 with PKM2 in proximity to K^305^ can be captured via TRAP from complex cell lysate, in agreement with crystallography^23^ (**Fig. 1i**) and TRAP analysis of recombinant PKM2 with TEPP-46 (**Extended Data Fig. 2a-b**). Moreover, this interaction is identified by TRAP across tested models including A549 lung carcinoma cell line and *E. coli* (**Fig. 1j, Extended Data Fig. 2c-d**), which verifies the unbiased nature of TRAP towards species for target identification. Then, we developed multiplexed-TRAP to further increase the throughput, which requires division of target cell lysates into aliquots followed by incubation with ligands of interest and solvent, separately (**Fig. 1k)**. After protein-level labeling, the proteome was digested, labeled with multiplexing reagents such as 6-plex Tandem Mass Tags (TMT), and fractionated followed by MS^3^ quantitative analysis. As expected, multiplexed-TRAP analysis also identified K^305^ of PKM2 that displays significantly reduced labeling occupation following TEPP-46 administration (**Fig. 1l-m**). Moreover, the accessibility at this responsive site decreases in a dose-dependent manner (**Extended Data Fig. 2e-f**). In contrast, the non-TRP harboring ^475^K that has negligible involvement in TEPP-46-PKM2 interaction exhibits constant accessibility (**Fig. 1n, Extended Data Fig. 2g**). Altogether, TRAP is poised to be a proteome-wide tool of identifying targetome and pinpointing interacting regions with no need of modification made to the original ligand.

### A Landscape of Glycolytic Targetome in Human Cancer Cells

Glycolytic metabolites are often defined as bioenergetics and biosynthetic precursors for cancer cells, albeit with their non-metabolic roles remaining encrypted. With the benchmarked TRAP approach, we sought to map glycolytic metabolites targetome in cancer cells and elucidate new sensory mechanisms and functional machinery of glycolysis. TRAP analysis of 10 glycolytic metabolites (**Fig. 2a**) was carried out in two batches using 6-plex TMT in HCT116 cells (**Fig. 2b**). In total, 160,459 peptides mapping to 6,926 unique proteins were identified, and 159,823 peptides belonging to 6,913 proteins were quantified. We first assessed the coverage of lysine-centric labels used in this study, and noted ~57.42% of lysine residues carrying the designated modification (**Fig. 2c, Extended Data Fig. 3a**). Consistently, the labeled fraction for quantified protein reaches ~57.34% (**Extended Data Fig. 3b-c**). Besides the relatively wide coverage^24^, distribution plot of the labeled lysine residues further shows the labeling reaction exploited by TRAP is not preferential towards any secondary structure elements (**Fig. 2d, Extended Data Fig. 3d**). The unbiased and reactive nature of lysine-centric labeling thus guarantees the discovery coverage of glycolytic metabolite targetome via TRAP. Overall, we initially identified 913 proteins as glycolytic metabolite target candidates and 2,487 interactions in HCT116 cells (**Fig. 2e, Extended Data Fig. 3e, Table S1**). For a total of 140 previously known metabolite-target interactions for *homo sapiens* documented in BRENDA that are quantified in this study, TRAP identified 30 known enzyme-metabolite interactions involving 19 proteins **(Extended Data Table S2**) and delivered a true Positive Rate comparable with previous stability-based approach^8^, which is expected to be further enhanced by overcoming proteomics undersampling (**Extended Data Fig. 3f**) and increasing the examined cell lines.

**Figure 2.**
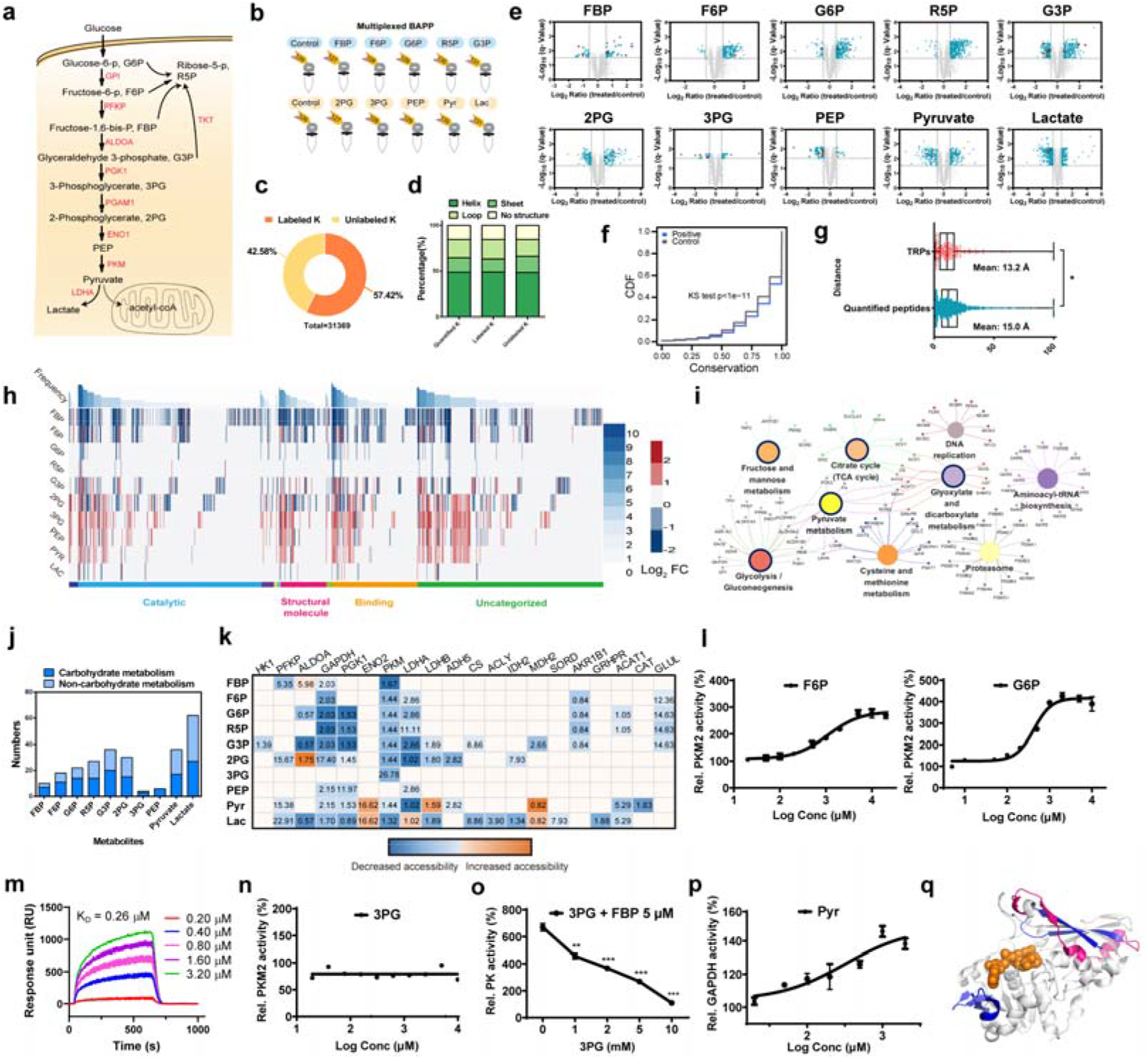
TRAP maps a global glycolytic targetome in cancer cells. **(a)** Glycolysis pathway in tumor cells. **(b)** A multiplexed workflow that efficiently identifies targetome for the given glycolytic metabolites in two batches of multiplexed-TRAP experiments. **(c)** Broad labeling coverage of reactive lysines achieved by the formaldehyde labeling-based TRAP approach. **(d)** Labeling preference analysis for high-order structures probed by TRAP. **(e)** Volcano plots of the glycolytic metabolites targetome mapped via TRAP. Proteins displaying significantly altered accessibility change following glycolytic metabolite incubation with the significance cutoff of q<0.03 and the TRAP ratio (treated/control) cutoff of >1.5 or <0.67 are color-coded in cyan with the previously known interactions highlighted in red. **(f)** Lysine conversation analysis of quantified peptides *vs.* TRPs of the assayed glycolytic metabolites. Statistical significance was evaluated by the Kolmogorov-Smirnov (KS) test. **(g)** Distance from the lysines contained in TRPs vs. Lysines of all quantified peptides to the functional sites of enzymes from the TRAP-identified targetome. **(h)** Gene Ontology (GO) molecular function (MF) classification of the identified glycolytic targetome showed enrichment in enzymes. **(i)** KEGG pathway annotation network analysis of catalytic proteins from the TRAP-identified glycolytic targetome. Pathways with p value <0.001 (by two side hypergeometric test) and the inclusion of at least three genes are summarized, with notable enrichment in the carbohydrate metabolism pathway (black circle). **(j)** Fraction of proteins involved in carbohydrate metabolism from the TRAP-identified enzymatic targets. **(k)** Accessibility changes of carbohydrate metabolism enzymes induced by glycolytic metabolite binding. Numbers in the boxes show the minimal Euclidean distances from the TRPs of the identified targets involved in the carbohydrate metabolism pathway to the closest atoms of substrates. Corresponding TRAP Ratio_Metabolite/Control_ is color-coded in the heatmap. **(l)** Enzymatic assay showed F6P and G6P as PKM2 activators. **(m)** SPR validated the engagement of human recombinant PKM2 with 3PG. **(n)** 3PG exerted negligible effect on PKM2 activation. **(o)** Co-administration of 3PG with FBP partially abrogated FBP-induced PKM2 activation. All comparisons were made between the PKM2 activity obtained from the co-administered group and that from the FBP treatment group. **(p)** GAPDH enzymatic activity assay showed pyruvate as an activator. **(q)** TRAP analysis suggests potential allosteric sites of GAPDH (PDB, 1u8f) since the TRPs of pyruvate include those within the active site (shown in blue) as well as those outside the active site with a minimum Euclidean distance of 21.54 Å (shown in purple). For (l) and (n)-(p), data represent mean ± SEM (n=3). *p < 0.05, **p < 0.01, ***p < 0.001, Student’s t-test.

Among the specific set of lysines in TRPs, we explored their functional ramifications and found the responsive sites are more conserved across organisms than all the quantified lysines (**Fig. 2f**). Consistently, since functional analysis of the identified glycolytic targetome suggests the enrichment in the category of catalytic proteins, we analyzed the lysines in the TRPs of these enzymes and found these residues are located more closely to the functional sites of given enzymes compared to those of quantified peptides, reiterating potential functionality of these glycolytic metabolites binding-responsive lysines (**Fig. 2g-h, Extended Data Fig. 4a**). Considering the fact that multiple glycolytic intermediates are well-recognized enzymatic ligands/substrates^8^, together we asked whether these findings suggest the assayed glycolytic metabolites are likely to be novel activity mediators of the identified enzymes in the target pool.

Since the enzymatic targets are further grouped to the carbohydrate metabolism pathway at highest frequency via KEGG pathway annotation network analysis (**Fig. 2i-j, Extended Data Fig. 4b, Table S3**), we focused on these carbohydrate metabolism enzymes and examined whether the TRAP-identified glycolytic metabolites interactions tend to influence the occupancy of active sites and concomitantly their activities. We define the boundary of an active site detectable by TRAP as 5.69 Å based on the Euclidean distances measured between the detected TRPs from known enzymatic targets that use these examined metabolites as substrates and the corresponding active sites (**Extended Data Fig. 4c, Table S4**). Therefore, when the distance from TRPs of given metabolites to an enzymatic active site falls within 5.69 Å, altered enzymatic activities are expected. We chose PKM2 and GAPDH, the two promiscuous glycolytic metabolites-binding partners identified by TRAP to test the proposed regulatory model (**Fig. 2k**). With the PKM2 activity assay, we found novel enzymatic activators F6P and G6P besides the classic activator FBP (**Fig. 2l, Extended Data Fig. 4d**), which is in line with their pronouncedly decreased accessibility in active site (**Fig. 2k, Extended Data Fig. 4e**). Interestingly, we noted 3PG as a PKM2 ligand, validated by surface plasmon resonance (SPR) (**Fig. 2m**), that harbors a TRP located distantly from the active site (~26.78 Å) among the examined glycolytic metabolites. In agreement, 3PG exerts negligible effect on PKM2 activity (**Fig. 2n**). Intriguingly, a K^433^-contaning TRP of 3PG is shared by FBP, indicating 3PG may potentially compete with FBP for binding to this region (**Extended Data Fig. 4f**). This inference was substantiated by the counteracted PKM2 activation upon co-administration of 3PG with FBP (**Fig. 2o**). Thus, 3PG might be defined as a partial agonist of PKM2, which hardly alters the enzymatic activity itself yet inhibits the FBP-induced activation. This finding suggests a complex and intrinsic feedback mechanism of cancer cells in precisely tuning cancer metabolism via the node of PKM2. In addition to identifying additional mediators for PKM2, the GAPDH enzymatic assay against the discovered ligand candidates surprisingly revealed that pyruvate can bind to and activate GAPDH (**Fig. 2p, Extended Data Fig. 4g**). The altered accessibility of lysines located remotely from the active site signifies a plausible allosteric mechanism heretofore unknown for GAPDH (**Fig. 2q, Extended Data Fig. 4h**). This allows design of novel chemicals targeting this ubiquitous and multi-functional protein that is intimately associated with various pathophysiology including cancer, immune diseases and neurodegeneration^25^.

### Uncover Multi-regulatory Modalities of Glycolytic Metabolites in Cancer Cells

Intriguingly, besides metabolic enzymes, targetome mining allows us to uncover noncanoical targets and thereby new functional modalities of glycolytic metabolites in cancer cells. Firstly, we found proteins ascribed to transcriptional factor activity by the Human Transcription Factor Database^26^ are identified via TRAP as glycolytic targets (**Extended Data Fig. 4a**), among which unligandable proteins such as TRIM28 are assigned as binding partners for the assayed glycolytic metabolites (**Fig. 3a**). Remarkably, TRIM28 is a universal nuclear co-repressor for Kruppel-associated box zinc finger proteins, and a SUMO and ubiquitin E3 ligase^27^. Its association with tumorigenesis is reported by mechanistic studies^27^ and supported by consistent upregulation in multiple cancer types and negative association with overall survival (**Extended Data Fig. 5a-b**). Among the TRAP-identified ligands, lactate changed the chemical accessibility of TRIM28 mostly, which indicates their stronger interactions compared to other assayed glycolytic ligands. We confirmed the proposed lactate-TRIM28 interaction by thermal shift assay in cell lysates (**Extended Data Fig. 5c**) and SPR (**Fig. 3b**). Subsequent TRAP analysis of cellular and recombinant TRIM28 both led us to note the lactate TRPs located in the coiled-coil (CC) domain of TRIM28 (**Fig. 3c-d, Extended Data Fig. 5d-e**), through which TRIM28 directly interacts with MDM2 and cooperates to promote p53 degradation^27^. Therefore, we wondered whether the accumulation of lactate in cancer cells is conducive to influence p53 levels dependent on TRIM28. Indeed, we treated HCT116 cells with doxorubicin (Dox) to induce p53 upregulation and found lactate repressed Dox-induced p53 level, whereas TRIM28 knockdown *per se* upregulated p53 and further abrogated the dampened p53 level conferred by lactate (**Fig. 3e, Extended Data Fig. 5f**). In agreement, intracellular deprivation of lactate by administering the LDHA inhibitor GNE140 increased p53 level while lactate replenishment partially revoked the increase. As expected, in siTRIM28 knockdown cells both effects were abolished (**Fig. 3f**). Together, these results indicate that the discovery of lactate-TRIM28 interaction and its influence on p53 degradation sheds light on a non-metabolic perspective regarding why lactate is exceedingly produced in cancer cells (**Fig. 3g**). Besides TRIM28, we also noticed lactate affected chemical accessibility and hence likely the conformations and functions of other transcriptional proteins that execute cancer malignant phenotypes exemplified by YBX1, a core regulator of MEK/ERK signaling-dependent gene expression signatures^28^, and NPM1, a multifunctional phosphoprotein implicated in tumorigenesis and immune escape^29^. Together with recent discovery of lactate as an epigenetic regulator by lactylation, our identification of lactate as a transcriptional regulator sheds light in understanding why the Warburg effect, characterized with lactate overproduction, is fundamental for cancer development and progression.

**Figure 3.**
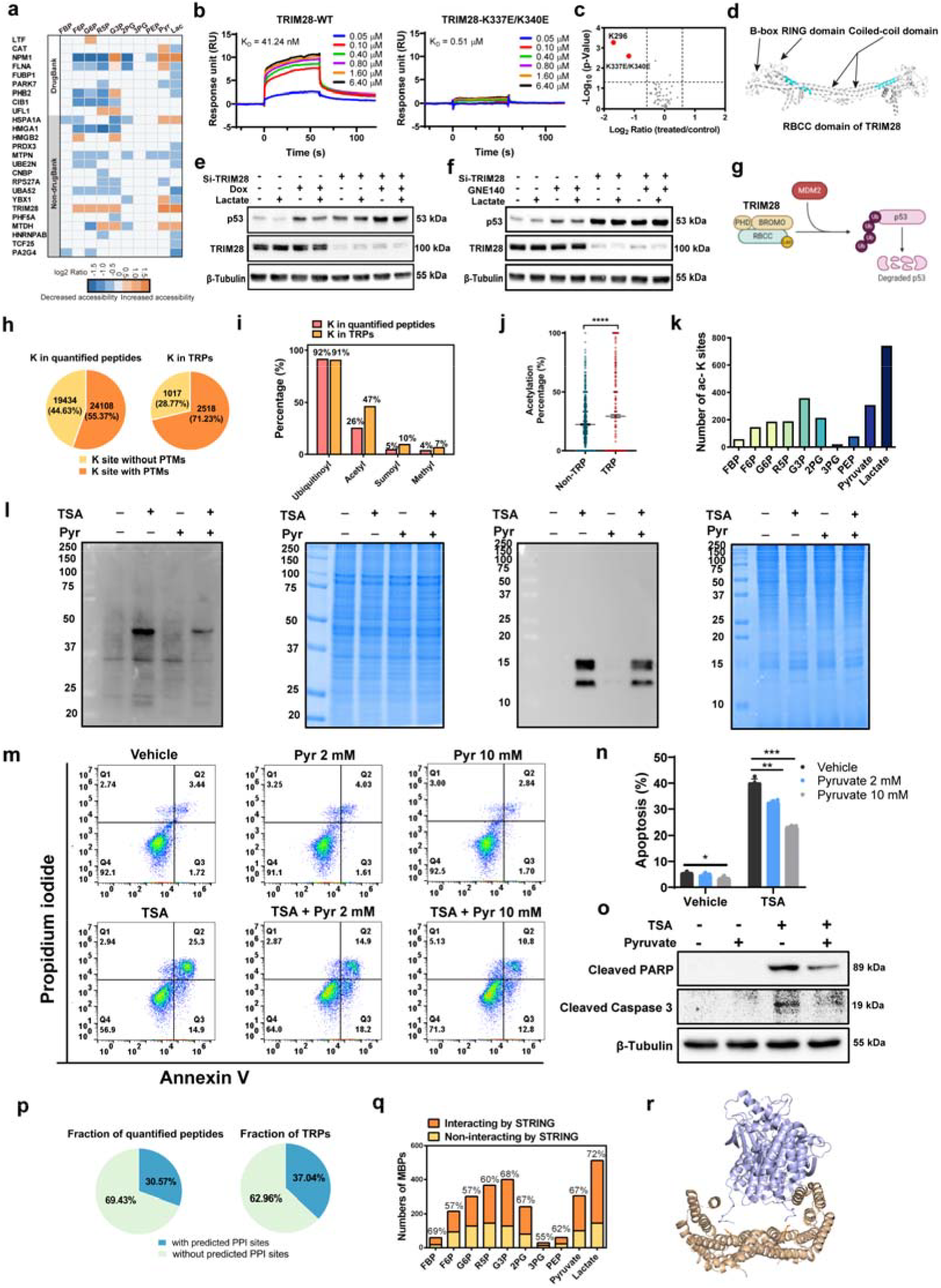
TRAP uncovers novel functional modalities of glycolytic intermediates. **(a)** Proteins of transcriptional regulation activity that exhibit changed accessibility following glycolytic metabolites administration. They are further classified by whether they have retrievable ligands from DrugBank. **(b)** Binding affinities of lactate to the wide type (WT) and the K337E/K340E double mutant TRIM28 measured by SPR. **(c)** TRAP analysis of human recombinant TRIM28 highlights the lysine residues that undergo significant accessibility changes in response to lactate binding. **(d)** X-ray structure shows the lactate-TRPs in TRIM28 is located in the Coiled Coil domain of TRIM28 (PDB: 6H3A) through which TRIM28 cooperates with MDM2 to promote p53 degradation. The identified TRP covering 328-347 is color-coded by cyan. **(e)** Abundance levels of p53 in control knockdown and TRIM28 knockdown cells following treatment with doxorubicin (Dox, 0.1 μM), lactate (10 mM) and both. **(f)** Replenishing lactate (10 mM) partially reversed the increase of p53 induced by GNE140 (10 μM), which was abrogated upon TRIM28 silencing. **(g)** Illustration of how lactate mediates TRIM28-mediated p53 degradation. **(h)** Fraction of lysines in quantified peptides *vs.* TRPs of the glycolytic targetome with annotation of post-translational modifications (PTMs) based on iPTMnet. **(i)** PTM category analysis of lysines in quantified peptides *vs.* TRPs of the glycolytic targetome. **(j)** Acetylation occupancy of lysines in non-TRPs *vs.* TRPs retrieved from identical glycolytic target proteins. ****p < 0.0001 by paired t-test. **(k)** Number of acetylated lysines in TRPs of glycolytic targetome based on documentation in iPTMnet for each assayed glycolytic metabolite. **(l)** Pyruvate (2 mM) administration reduced pan-acetylation level upregulation induced by trichostatin A (TSA, 0.5 μM) in HTC116 cells. **(m)** Pyruvate administration reduced the apoptotic rate of HCT116 cells induced by TSA (0.5 μM) according to AV-FITC/PI staining by flow cytometry. **(n)** Statistic results of cell apoptosis by flow cytometry. Data represent mean ± SEM (n=3). *p < 0.05, **p < 0.01, ***p < 0.001 by Student’s t-test. **(o)** Immunoblotting of cleaved PARP and cleaved caspase 3 expression in HCT116 cells following treatment with TSA (0.5 μM), pyruvate (2 mM) and both. β-Tubulin was used as the loading control. **(p)** Fraction of quantified peptides *vs.* TRPs of glycolytic metabolites with predicted protein-protein interaction events by prePPI. **(q)** STRING analysis of protein-protein interaction partners among the TRAP-detected glycolytic targetome. **(r)** X-ray structure of 14-3-3ζ (wheat, cartoon) and B-RAF (blue, cartoon) shows the pyruvate/lactate-binding responsive site K^49^ (orange, stick) in 14-3-3ζ is located at the protein-protein interacting site (PDB, 6U2H).

Aside from transcriptional regulation that directly controls downstream genes and protein abundance, PTM serves as a core mechanism of switching protein conformation and activity. Driven by the question of whether glycolytic metabolites binding affects proteome PTM levels, we retrieved PTM evidence for lysines in TRPs and noted these regions are more likely to carry PTMs compared to the total quantified proteome (**Fig. 3h**). It is noteworthy that acetylation was enriched for ~2 folds for lysines of TRPs (**Fig. 3i**). To exclude the possibility that differences in the frequency of acetylated lysines result from heightened percentage of acetylation in assigned target proteins, we compared the frequency of lysine acetylation for lysines of TRPs with those of non-TRPs yet from identical proteins, and also noted enriched acetylation (**Fig. 3j**). GO analysis shows these proteins do not distribute in exclusive subcellular location or specific class compared to the quantified proteome (**Extended Data Fig S6a-b**). Therefore, we conjecture a prevalent acetylation-based regulatory mechanism for glycolytic metabolites achieved by engagement-hindered surface exposure to acetylation writers. Since pyruvate and lactate are top-ranked metabolites that bear acetylation sites in their TRPs (**Fig. 3k**), we analyzed the impact of pyruvate and lactate treatment on proteome pan-acetylation levels by immunoblotting in HCT116 cells, and found that they can both partially abrogated acetylation induced by trichostatin A (TSA) treatment, a mammalian histone deacetylase (HDAC) inhibitor^30^, with the most marked decrease observed for histone (**Fig. 3l, Extended Data Fig. 6c**). Projecting forward, we confirmed the influence of pyruvate on histone acetylation functionally affected the pharmacological phenotypes correlated with TSA exemplified by the mitigated apoptotic rate (**Fig. 3m-n**) and reduced PARP and cleaved caspase-3 levels (**Fig. 3o**). Besides histone, we found pyruvate also influence accessibility of lysines from a myriad of non-histone target proteins. A notable example is K^215^ from the GAPDH TRP that displays the most dramatic accessibility change in response to pyruvate administration (**Extended Data Fig. 4h**). Collectively, the impact of glycolytic metabolites such as pyruvate on acetylation of diverse proteins expands our understanding of how glycolysis may be intimately linked with posttranslational regulation.

Lastly, protein-protein interactions (PPI) have been verified as essential players in signal transduction while targeting PPI is a promising but largely unexplored area for drug discovery. It is also unclear about whether glycolytic intermediates regulate PPI in cancer cells. Through PPI analysis of the glycolytic targetome, we observed a tempered enrichment for lysines of TRPs compared to those of the quantified proteome as known or predicted PPI sites by prePPI, implying metabolites binding can mediate the equilibrium of protein complexes assembly (**Fig. 3p**). Accordingly, STRING analysis shows a majority of targets, ranging from 55-72%, are experimentally validated interacting partners with high confidence (interaction score> 0.9) (**Fig. 3q**). An exemplary case is the finding that lactate and pyruvate can both change the accessibility of S28-R55 in the adapter protein 14-3-3ζ, in which K^49^ is known to coordinate with the phosphorylation moiety in both B-Raf and C-RAF kinases according to crystallography^31,32^ (**Fig. 3r**). Thus, the increased flux of pyruvate and lactate through activating glycolysis may coordinate with the 14-3-3ζ-RAF complex assembly (**Extended Data Fig S6. d-e**) and thereby modulate the central Ras-RAF-MEK-ERK pathway^31,32^. Altogether, TRAP analysis allows us to uncover versatile regulatory modalities for glycolytic metabolites including transcriptional control, PTM rewiring and protein assemblies modulation, revealing a global targetome that can be intrinsically directed by glycolytic dynamics in cancer cells.

### Harnessing the Glycolytic Metabolite Targetome for Anti-cancer Translational Research

Since glycolysis is a hallmark of cancer, the contributing glycolytic enzymes and related signaling nodes have been exploited as possible targets for cancer therapy^33^. Our findings together with previous studies support that glycolytic intermediates are important mediators in tuning cancer metabolism and signaling transduction, and thus we reasoned that glycolytic metabolites themselves might be explored for their potentials as leading compounds for drug design. To this end, we employed NaF and GNE140 as representative glycolysis inhibitors, and examined whether the accumulated metabolites upon the intervention are functional and thus constitute a fertile repertoire of leading compound template for drug discovery. Expectedly, we noted suppressed proliferation following NaF and GNE140 administration^34,35^ (**Extended Data Fig. 7a)**. This is accompanied with pronouncedly altered abundance of glycolytic intermediates (**Fig. 4a**, **Extended Data Fig. 7b**). Screening of the increased glycolytic metabolites indicated that G3P most significantly suppressed HCT116 cell growth (**Extended Data Fig. 7c**). We thus treated HCT116 cells with G3P (**Fig. 4b**) and found it induced dose-dependent inhibition on HCT116 cell growth (**Fig. 4c**) as well as on HT29 cells (**Extended Data Fig. 7d**). The inhibitory effect of G3P on cancer cell proliferation was further demonstrated by colony formation assay (**Fig. 4d**). These findings led us to propose the query of G3P as a leading compound for anti-cancer drug discovery and, more importantly, its interacting proteins hold promise to uncover a new panel of therapeutic targets that combinatorially control cancer cell growth.

**Figure 4.**
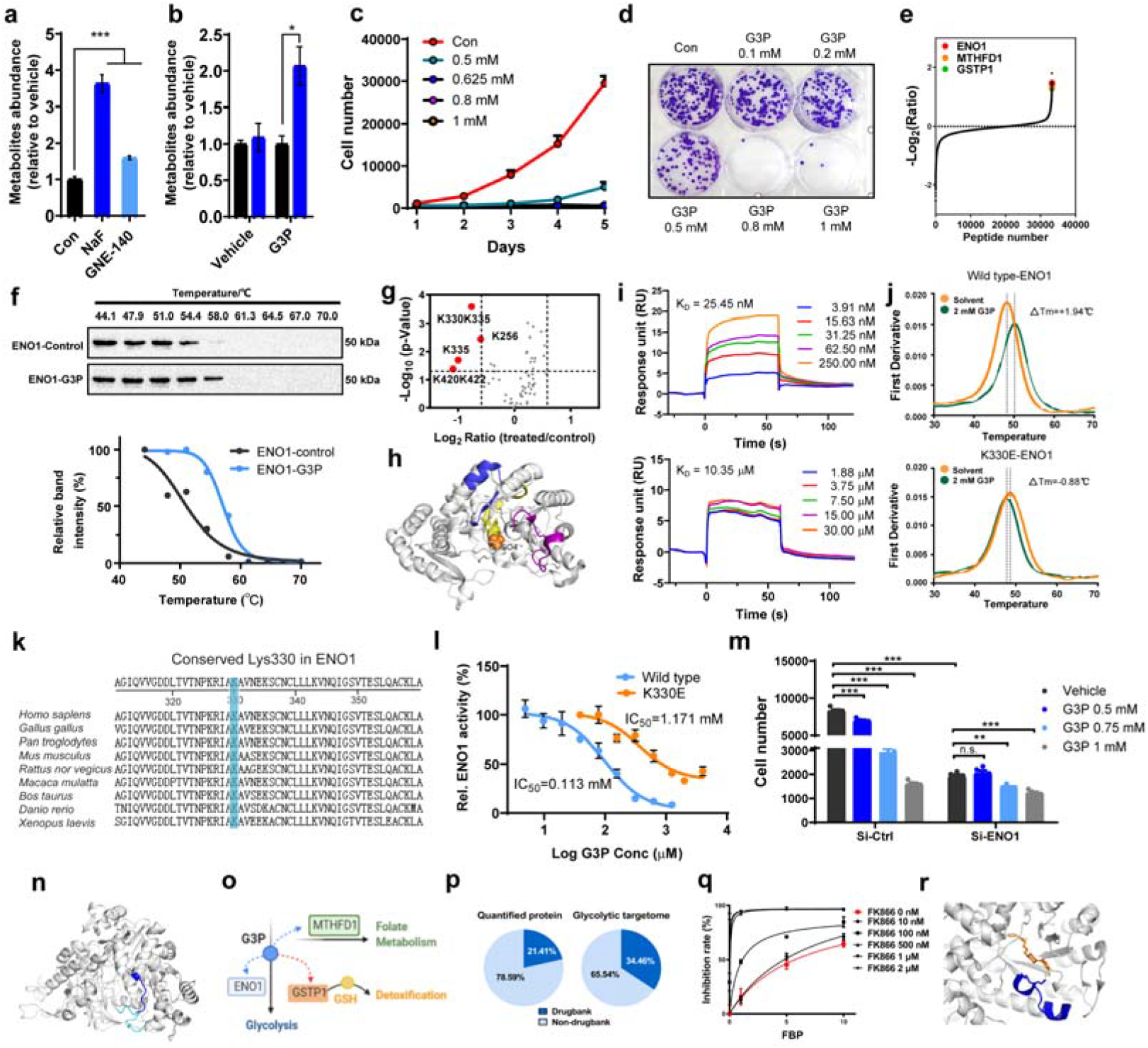
G3P is an intrinsic proliferation-suppressive metabolite via targeting ENO1 and MTHFD1. **(a)** Intracellular concentrations of G3P in HCT116 cells treated with solvent control, NaF (3 mM) and GNE140 (10 μM) for 24 hr determined by targeted LC/MS-MS (n = 3). **(b)** Accumulative G3P upon direct supplementation of 0.5 mM G3P for 4 hr in HCT116 cells (n = 3). **(c)** Growth curve of HCT116 cells under G3P administration (n = 5). **(d)** Colony formation assay validates G3P-induced suppressive proliferation in HCT116 cells. **(e)** S-plot of TRPs shows top-ranked target proteins of G3P that displayed marked accessibility changes following G3P administration. **(f)** Thermal shift assay validated the ENO1-G3P engagement in HCT116 cell lysates. **(g)** Volcano plot of the TRAP-identified G3P TRPs in recombinant ENO1. Accessibility of the TRP carrying Lys330/Lys335 exhibited the most dramatic change after G3P incubation. **(h)** The ENO1 TRPs of G3P identified by TRAP are highlighted (PDB: 3B97). TRPs spanning residue 328-343, 407-426 and 254-269 are color-coded in blue, yellow and purple. **(i)** Binding affinities of G3P to the wide type (WT) and the K330E mutant ENO1 measured by SPR. **(j)** Stability changes of the WT and K330E mutant ENO1 induced by G3P incubation examined by nanoDSF. **(k)** Conservation analysis of K330 of ENO1 among several species. **(l)** Enzyme activity assay showed G3P inhibited the activity of WT ENO1 yet weakly to the K335E mutant ENO1 (n = 3). **(m)** Proliferation of the control and ENO1 knockdown cells under G3P treatment (n = 5). **(n)** Homology modeling of MTHFD1 indicates the TRP of G3P is located in proximity to the ATP binding site and thus the interaction likely interferes with the 10-formyl-THF synthetase reaction. **(o)** Multi-target regulatory network of G3P uncovered by TRAP. **(p)** DrugBank and non-DrugBank fraction of the identified glycolytic targetome. **(q)** NAMPT enzymatic assay suggests synergistic inhibition by FBP with FK866. **(r)** TRAP analysis reveals the TRP (blue, cartoon) of FBP located in proximity to FK866 (orange, stick) displayed reduced accessibility in response to FBP (n = 3), suggesting a FBP binding-promoted interaction between FK866 and NAMPT (PDB: 2GVJ). For (a)-(c) and (l)-(m), data represent mean ± SEM. For (a, b, m), *p < 0.05, **p < 0.01, ***p < 0.001, n.s. non-significant, Student’s t-test.

We thus plotted the TRAP R_G3P/Control_ for G3P, and the resultant S-plot of TRPs suggests α-enolase (ENO1), methylenetetrahydrofolate dehydrogenase (MTHFD1) and glutathione S-transferase Pi 1 (GSTP1) as top-ranked G3P-binding targets (**Fig. 4e**). We confirmed the three top-ranked target candidates as the binding partners of G3P, since they all become stabilized by G3P in the tested temperature range compared to the vehicle (**Fig. 4f, Extended Data Fig. 7e-f**). An iso-thermal dose-response experiment further validated the G3P-induced stabilizing effect (**Extended Data Fig. 7g**). Further, with ENO1 displaying the most pronounced accessibility change (R_G3P/control_ =0.36), we first sought to dissect the connection of G3P-ENO1 binding with the growth-inhibitory function of G3P. We applied TRAP to screen the possible binding sites of G3P to recombinant ENO1, and noted a TRP containing K^330^ showing significantly reduced accessibility following G3P incubation (**Fig. 4g-h, Extended Data Fig. 7h**). Thus, we examined the binding affinities of G3P to the wild-type (WT) ENO1 and the K330E ENO1, and confirmed the point mutation at K^330^ weakened the G3P-ENO1 interaction by changing the K_D_ from 25.45 nM to 10.35 μM (**Fig. 4i**). The interaction between G3P with the K^330^-proximal region was further validated via differential scanning fluorimetry (DSF), since the G3P-mediated protection of ENO1 against heat-induced denaturation was abrogated using K330E ENO1 (**Fig. 4j**). Conservation analysis of ENO1 showed that the K^330^ are highly conserved, which suggests the importance of K^330^ in preserving the function of ENO1 in evolutionarily divergent species (**Fig. 4k**). Therefore, we tested G3P on ENO1 activity and determined G3P as an inhibitor, whereas the K330E mutation abrogated the inhibition of G3P on ENO1 (**Fig. 4l**). Then, we conjectured that ENO1 mediates G3P-induced proliferation arrest. To validate this hypothesis, we used siRNA to silence ENO1 (**Extended Data Fig. 7i**). ENO1 knockdown *per se* significantly retarded the proliferation of HCT116 cells, and largely abolished the proliferation-arrest effect of G3P (**Fig. 4m**). Together, these results support that G3P-mediated inhibited growth of HCT116 is at least partially dependent on ENO1.

Subsequently, we wondered whether the interactions of MTHFD1 and GSTP1 with G3P also mediate the G3P-induced growth suppression of HCT116 cells. For MTHFD1, TRAP analysis reveals that G3P affects the accessibility of the region proximal to the ATP-binding pocket in the 10-formyl-THF synthetase domain of MTHFD1 by homology modeling (**Fig. 4n**), potentially leading to impaired 10-formyltetrahydrofolate synthesis mediated by MTHFD1^40^. Expectedly, MTHFD1 knockdown also markedly inhibited HCT116 cell growth and partially abrogated the G3P-induced growth arrest (**Extended Data Fig. 7i-j**). Conversely, although TRAP analysis suggests G3P binding influences the accessibility of the substrate-binding domain of GSTP1 (**Extended Data Fig. 7k**), which possibly interferes with glutathione conjugation and hence the detoxification capacity of cells^37^, GSTP1 knockdown did not significantly inhibit HCT116 cells growth and hardly impacted the G3P-mediated growth suppression under the examined condition (**Extended Data Fig. 7i-j**). Collectively, we identified non-metabolic regulation mechanisms for G3P. Besides establishing a negative-feedback loop between glycolysis and G3P, our findings also reveal crosstalk capacity of glycolysis with pathways involving folate metabolism and possibly redox balance, and reiterate an intricate glycolytic metabolite-protein network that underlies complex cancer cell machinery (**Fig. 4o**). We also anticipate G3P serves as a template for designing new anti-cancer therapeutics targeting ENO1 and MTHFD1.

Glycolysis is itself an attractive and promising area for anti-cancer drug discovery and moreover it has been well demonstrated that cancer metabolism is closely associated with clinical responses to cancer therapeutics. Since screening of the identified glycolytic targetome shows they are enriched as druggable proteins compared to the quantified proteome (**Fig. 4p, Extended Data Fig. 8a)**, we are of particular interest in exploring whether these metabolites may interact with cancer therapeutics, and ultimately modulate the acquired pharmacological outcomes by direct engagement. Specifically, we found FBP administration dose-responsively inhibited nicotinamide phosphoribosyltransferase (NAMPT), an essential protein for NAD^+^ biosynthesis in cancer cells^38^, and synergized with FK866, the NAMPT inhibitor under clinical investigation (**Fig. 4q)**. Mechanistically, TRAP shows FBP binding reduced accessibility in proximity to the FK866 binding site (~3.42 Å), and explains the enhanced affinity of FK866 to NAMPT by FBP treatment (**Fig. 4r**). Additionally, pyruvate, a newly identified GAPDH activator (**Fig. 2p**), significantly compromised dimethyl fumarate (DMF)-mediated inhibition of GAPDH^25^ **(Extended Data Fig. 8b-c**). These findings substantiate an important concern that the metabolic context and variation may represent a key factor in determining individual variations in response to cancer therapeutics, supporting that glycolysis can be harnessed as adjuvants for currently available cancer therapeutic options.

## DISCUSSION

With the accessibility profiling-based TRAP approach, we mapped a global glycolytic targetome from cancer cells, and pinpointed the proximal regions where the interactions occur. The depicted topology of metabolite-target interactions enables us to extrapolate functional principles regarding how glycolytic metabolites diversely and comprehensively regulate the binding proteins and thereby shedding lights on understanding how they are engaged in coupling metabolism to tumor signaling in support of tumor growth and metastasis. The glycolytic targetome identified in our study supports the recently revisited functions of glycolytic metabolites-not just as metabolic fuels and biosynthetic building blocks, but also as central mediators that diversely regulate carbohydrate metabolism, nuclear transcription, protein PTM dynamics and macromolecular assemblies in cancer cells.

In particular, our findings of lactate as a ligand of an orphan transcriptional factor TRIM28 facilitating p53 degradation and multiple metabolites attenuating histone acetylation and hence TSA-induced apoptosis reflect the survival advantages gained by cancer cells via the hyperactivated glycolysis. The p53 protein, a fundamental sensor and checkpoint of DNA damage and metabolic state, serves as a master regulator of tumor development and progression via transcriptional control of multiple genes involved in cell fate decision and metabolism^39^. However, it remains largely unexplored regarding how glycolysis reciprocally regulates p53. Our identification of lactate as a negative regulator of p53 proteostasis thus unravels a mechanistic connection in how p53 signaling is precisely intertwined with glycolysis, partially revealing a complex communication network orchestrated by cancer cells, and suggests a therapeutic strategy in strengthening p53 response.

Intriguingly, besides pro-tumoral signaling metabolites, we also identified an intrinsically bioactive intermediate G3P that effectively blocks the proliferation of colorectal cancer cells. Integration of biochemical experiments allows us to demonstrate ENO1 and MTHFD1 from the TRAP-assigned G3P targetome as essential nodes responsible for the G3P accumulation-mediated growth arrest. These results substantiate the existence of inter-pathway crosstalk between glycolysis and folate metabolism, and more excitingly, suggest G3P as an anticancer leading compound that targets both ENO1 and MTHFD1, for which warrants future research in validating the potential combinatorial and synergistic effects in combating tumor growth and progression. In fact, emerging evidences support that endogenous metabolites-inspired candidates design constitute a fertile resource for innovative drug development. This has been exemplified by α-ketoglutarate (αKG)^40^, itaconate^7^, and glutamine^41^, for whose targets are exploited to combat cancer and overcome tumor immune evasion by administering metabolites-derived mimetics or competitors. Another translational finding is that the identified glycolytic targetome consists of a wealth of druggable targets. This implies dynamic changes of given glycolytic ligands might alter the conformations of their interacting drug targets and hence affect drug binding affinity and occupancy. In this context, we believe that monitoring glycolytic biomarkers is essential in determining individual variations in response to cancer therapeutics and its manipulation represent therapeutic opportunities to improve precision and optimize clinical benefits.

In summary, the glycolytic targetome identified via TRAP in this study provides a new perspective to systematically explain how hyperactive glycolysis is fully utilized by cancer cells via finely tuning metabolism with cellular signals. Moreover, the previously unannotated glycolytic metabolites-binding proteins may lead to the discovery and validation of promising targets for anti-cancer therapeutics development, and likewise, the identification of ligands to proteins previously defined as unligandable can thus immediately inspire drug design that derives from the glycolytic ligands.

Together with recent chemoproteomics-empowered targetome mapping and regulatory mechanisms elucidated for multiple functional metabolites, we have realized isolated observation and recording of proteins and metabolites abundance and cellular/tissue distributions are insufficient, whereas the knowledge gap of metabolite-protein interactions constantly occurring in nature must be bridged-to obtain a holistic picture of the target landscape for endogenous metabolites especially in human cells/tissues, for which a program of Human Metabolite Targetome may be proposed, calls for collaborative efforts and global contributions.

## ACKNOWLEDGEMENTS

This research was supported by the National Natural Science Foundation of China (grants 81930109, 81720108032 and 91429308 to H.H., grants 81872838 to H.Y.), the Natural Science Foundation of Jiangsu Province (BK20180079), the Project for Major New Drug Innovation and Development (grants 2018ZX09711001-002-003 and 2018ZX09711002-001-004 to H.H., 2018YFD0901101 to H.Y.), Overseas Expertise Introduction Project for Discipline Innovation (grant G20582017001 to H.H.), Double-First Class Initiative Project (grants CPU2018GF09 to H.H., CPU2018GY09 to H.Y.), State Key Laboratory of Natural Medicines at China Pharmaceutical University (grants SKLNMZZCX201817 to H.H. and SKLNMZZCX201801 to G.W.), and Sanming Project of Medicine in Shenzhen (SZSM201801060). We thank Prof. Yibei Xiao from China Pharmaceutical University, Dr. Qing Yu from Harvard University and Dr. Baozhen Shan and Wenting Li from PEAKS Studio for useful discussions.

## AUTHOR CONTRIBUTIONS

H.H., H.Y. and G.W. conceived the project. Y.T., N.W., H.Y. and H.H. designed the experiments. Y.T., N.W., H.Y. and M.D. performed the proteomics experiments. Y.T. and C.S. performed the flow cytometry and western blotting experiments, Q.B. performed the SPR experiments, N.W. conducted the pan-acetylation immunoblotting experiment, W.L., K.Z. and S.C. carried out protein site-mutation and purification experiments, H. H. conducted the conservation analysis, H.S. performed the homology modeling analysis of MTHFD1, C.Y. assisted with partial proteomics data acquisition, N.W., Y.T., H.Y. and H. H. analyzed the experimental data. H.Y. and H.H. wrote the paper with input from co-authors.

## METHODS

### Cell culture and chemicals

All cells were all obtained from American Type Culture Collection and grown at 37 °C with 5% CO_2_. HCT116 (CCL-247) cells were cultured in McCoy’s 5A Medium, and HT29 (HTB-38) cells were cultured in Dulbecco’s Modified Eagle Medium (DMEM) purchased from Gibco (Grand Island, NY, USA). All medium were supplemented with 10% fetal bovine serum (FBS) (Biological Industries, Kibbutz, Israel), 100 unit/mL penicillin and 100 μg/mL streptomycin (Thermo Fisher Scientific, San Jose, CA, USA). LC-MS grade water and acetonitrile (ACN) were obtained from Merck (Darmstadt, Germany). All chemicals were purchased from Sigma Aldrich (St Louis, MO, USA) unless otherwise specified.

### TRAP workflow for purified/recombinant proteins

Model proteins including purified bovine RNase A and human recombinant PKM2 were incubated with their corresponding ligands or solvent for 1 hr at room temperature. The proteins were labeled with 2 μL of 0.5% CD_2_O and 10 μL of 10 mM Borane Pyridine Complex (BPC) (J&K Scientific Ltd., Beijing, China, cat. no. 121499) for 30 min, and the reaction was quenched by the addition of NH_4_HCO_3_ to a final concentration of 50 mM and incubation for 20 min. The resultant mixtures were filtered with 10 kDa molecular weight cut-off (Merck, Darmstadt, Germany) by centrifugation at 12,000 g for 15 min. Then, the enriched proteins were denatured by 8 M urea for 30 min followed by incubation with 5 mM dithiothreitol (DTT) at 56 °C for 30 min and subsequent incubation with iodoacetamide (IAM) at room temperature in dark for 30 min. DTT was added again to neutralize the excess IAM for another 10 min. The sample was then diluted with 25 mM NH_4_HCO_3_ to make the concentration of urea less than 1 M, and overnight digested by trypsin in an enzyme/protein ratio of 1:40 (wt/wt) at 37 °C. The digestion was quenched by formic acid (FA) followed by desalting with C_18_ Ziptip (Waters, Milford, MA, USA).

### TRAP workflow for human cancer cell line proteome

#### Cell lysates preparation

Cells were washed with cold phosphate buffered saline (PBS) for three times, scraped and centrifuged at 1,000 rpm for 5 min. The cell pellets were resuspended in mammalian protein extraction reagent (M-PER, Pierce/Thermo Fisher Scientific, cat. no. 78503) containing protease inhibitor (ApexBio Technology, Houston, TX, USA, cat. no. K1007) and phosphatase inhibitor (ApexBio Technology, cat. no. K1013), and lysed on ice for 30 min. The supernatants were collected by centrifugation at 18,000 g for 10 min. Protein concentration was determined by bicinchoninic acid (BCA) Protein Assay (Beyotime Biotechnology, Beijing, China, cat. no. P0011). The cell lysates were then diluted with M-PER lysis to 3 μg/μL and incubated with given metabolites or solvents for 1 hr at 25 °C. Specifically, the glycolytic metabolites were administered at dosages detailed as follows: D-Fructose 1,6-bisphosphate (FBP, 200 μM), D-fructose 6-phosphate (F6P, 100 μM), D-glucose 6-phosphate (G6P, 100 μM), D-ribose 5-phosphate (R5P, 30 μM), D/L-glyceraldehyde-3-phosphate (G3P, 30 μM), D-2-phosphoglycerate (2PG, 10 μM) from Toronto Research Chemicals, North York, Ontario, Canada. D-3-phosphoglycerate (3PG, 10 μM), phosphoenolpyruvate (PEP, 5 μM), pyruvate (Pyr, 300 μM), L-lactate (Lac, 2 mM). The dosages were set based on the intracellular concentrations of these metabolites as reported in literature^42,43^.

#### Proteome labeling and preparation for the label-free quantification-TRAP workflow

Cell lysates (150 μL, 3 μg/μL) were labeled by the addition of 6 μL of 1% CD_2_O and 90 μL of 10 mM BPC for 30 min. The reaction was quenched by incubating with 50 mM NH_4_HCO_3_ for 20 min. Then, methanol, chloroform and water were added to the labeled lysate in order according to a ratio of 4:1:3:1 by volume followed by centrifugation at 12,000 rpm for 10 min to precipitate the proteome. The flaky precipitate was washed twice with methanol and re-solubilized by 8 M urea in 25 mM ammonium bicarbonate solution. Proteins were then reduced by 10 mM DTT by incubation at 56 °C for 30 min, and alkylated by 40 mM IAM at 25 °C for 20 min in dark. Additional DTT was added and allowed to react with excess IAM at room temperature for 10 min. The mixture was added with 25 mM NH_4_HCO_3_ to dilute urea to 1 M followed by overnight digestion with sequencing-grade trypsin (Promega, Madison, WI, USA) at an enzyme/protein ratio of 1:40 (wt/wt) at 37 °C. The digestion was quenched by FA addition to pH=3. The mixture was desalted with C_18_ SepPak cartridges (Waters) and evaporated to dryness with vacuum centrifuge (Thermo Fisher Scientific). Samples were stored at −80 °C prior to analysis.

#### Sample preparation for TMT-based multiplexed TRAP

Cell lysates were labeled and precipitated using the same protocol as the label-free quantification-TRAP workflow. The precipitate was resuspended in 8 M urea solution (in 50 mM Tris HCl containing 10 mM EDTA, pH 8.0) followed by DTT reduction and IAM alkylation. To this was added LysC (Signalchem, L585-31N-05) in an enzyme/substrate ratio of 1:400 (wt/wt), followed by incubation at 25 °C for 4 h. The mixture was diluted with 25 mM NH_4_HCO_3_ so that the final concentration of urea is less than 2 M, and then sequencing grade trypsin was added in an enzyme/substrate ratio of 1:50 for overnight digestion at 37 °C. After quenching by FA, the mixture was desalted with reversed phase SPE cartridges (Waters) and dried with vacuum centrifuge. The sample was resuspended in 300 μL of 50 mM HEPES (pH 8.5) with the concentration of peptides determined by Pierce quantitative colorimetric peptide assay (Rockford, IL, USA). A 60 μg peptide aliquot of each sample was labeled with TMT reagent for 1.5 hr according to the manufacturer’s instructions. The reaction was quenched by incubating with 10 μL 5% hydroxylamine for 15 min. Aliquots labeled with the 6-plex TMT reagents were combined and acidified with FA followed by evaporation to dryness. The labeled proteome is then desalted again by SPE cartridges and lyophilized. The proteome was dissolved in HPLC phase A buffer (10 mM ammonium formate containing 5% acetonitrile) and then injected into the sample loop of the UPLC system (Acuity, Waters,) for fractionation. Phase B consists of ACN with 20% 10 mM ammonium formate aqueous buffer. The gradient was set as follows: 0-5 min, 1% B; 5-79 min, 1-50% B; 79-81 min, 50-100% B; 81-98 min, 100% B; 98-100 min, 100-1% B; 100-120 min, 1% B. The effluent was collected every 1.5 min. Every 12 fractions were set as a cycle, and each fraction was combined with the fractions collected in the following cycles. The lyophilized fractions were dissolved in 60 μL of 0.1% FA followed by desalting with C_18_ Ziptips and storage at −80 °C prior to analysis.

### Mass spectrometry

#### Intact mass measurement

RNase A (100 μM) and CDP/CTP (1 mM) was both dissolved in 25 mM ammonium acetate buffer and incubated for 30 min, and native MS measurement of the formed holo-complex was conducted on a TripleTOF 5600 system (SCIEX, Framingham, MA, USA) by direct infusion. The instrument was set to acquire over the *m/z* range of 100−2000 Da for TOF-MS scan.

For intact mass measurement of the dimethylated RNase A with and without ligand incubation, labeled RNase A was desalted by 3 kDa MWCO and analyzed on a C_4_ column (4.6×150 mm, 3 μm, 300 Å, Sepax Technologies, Newark, DE, USA) on an LC-30 HPLC system (Shimadzu, Kyoto, Japan). The mobile phase consisting of 0.1% FA in water (phase A) and 0.1% formic acid in ACN (phase B) was delivered at a flow rate of 0.4 mL/min using a 15 min gradient program. The eluent was then introduced via ESI ion source into the TripleTOF 5600 system ((AB Sciex, Framingham, MA, USA)) for mass measurement. Q-TOF analyzer was set to scan over the *m/z* range of 100-2000. The spectra were combined by summing across the chromatographic peak of labeled RNase A and deconvoluted using the SCIEX BioPharma View Software.

#### Label-free proteomic quantification

Data used for label-free quantification was acquired on a nanoACQUITY UPLC system coupled to SYNAPT G2-Si mass spectrometer (Waters). A C_18_ trapping column (Waters Acquity UPLC M-Class, 0.18×20 mm, 5 μm, 100 Å) and a HSS T3 analytical column (Waters Acquity UPLC M-Class, 75 μm×150 mm, 1.8 μm, 100 Å) were employed. Mobile phases A and B consist of 0.1% FA in water and 0.1% FA in ACN, respectively. A 60 min and 120 min length gradient of 1-40 % acetonitrile at a flow rate of 300 nL/min was used for separation of recombinant protein digests and cell lysates samples, respectively. MS scan range was set to *m/z* 350-1500 with a scan time of 0.2 s, and MS/MS scan range was set to *m/z* 50-2000 using data-dependent acquisition (DDA). The top 10 abundant precursors were subjected to MS/MS fragmentation with a ramp CE set between low energy (14-19 eV) and elevated energy (60–90 eV) using a scan time of 0.15 s per function.

#### TMT-based MS3-level multiplexed quantification

Data used for the multiplexed TRAP workflow was collected on an Orbitrap Fusion Lumos mass spectrometer equipped with an EASY-nano LC 1200 liquid chromatography system (Thermo Fisher Scientific). Mobile phase A consisting of 0.1% FA in water and B consisting of 0.1% FA in ACN-H_2_O (8:2 by volume) were delivered at a flow rate of 300 nL/min. The 75 μm capillary column was packed with 35 cm of Accucore 150 resin (2.6 μm, 150 Å, Thermo Fisher Scientific). Peptides were analyzed using a 150 min chromatography gradient from 0%-50% phase B during 5-79 min. For MS data acquisition, MS1 spectra were collected at the *m/z* range of 375-1500 at a resolution of 120,000 in the Orbitrap with a maximum injection time of 50 ms or a maximum automated gain control (AGC) value of 4e^5^. For MS2 acquisition, fragmentation was conducted by collision-induced dissociation with a normalized collision energy (NCE) at 35. MS2 spectra were collected at the mass range of 400-1200 in ion trap with a maximum AGC of 1e^4^ or a maximum injection time of 50 ms. For accurate quantification, MS3 were conducted for TMT reporter ion quantification by high energy collision-induced dissociation (HCD) with NCE at 65. The MS3 spectra were collected over the mass range of 100-500 at a resolution of 50,000 with the maximum injection time set at 10^5^ ms and AGC target value at 1e^5^.

### Proteomic data analysis and bioinformatics

#### Protein identification and quantification

The acquired label-free DDA data and TMT-MS3 data were searched against the *Homo sapiens* UniProt database (version 2018) using PEAKS Studio 8.5 (Bioinformatics Solutions Inc., Waterloo, Ontario, Canada). Due to the necessary lysine labeling step employed by TRAP, we allowed up to two missed cleavages and semi-specific tryptic digestion. Carboxyamidomethylation on cysteines (+57.02 Da) was selected as fixed modification, and methionine oxidation (+15.99), CD_2_O-mediated dimethylation (+32.06 Da) and mono-methylation (+16.03 Da) on lysines were set as variable modifications. For label-free quantification data, precursor mass tolerance was set to 20 ppm, and fragment mass tolerance was set to 0.1 Da. For TMT-based MS3 quantification data, precursor mass tolerance was set to 10 ppm, MS2 fragment mass tolerance was set to 0.6 Da, and MS3 fragment mass tolerance was set at 0.02 Da. The identified proteins were filtered with 1% FDR and the quantified proteins must include at least one unique peptide. Regarding label-free quantification, we set 50 ppm mass tolerance and 3 min retention time shift tolerance for peptide alignment.

#### Classification of quantified peptides and proteins

We assessed potential glycolytic targetome by removing proteins associated with keratin firstly from the pool of quantified proteins, and only considered the quantified peptides which belong to the following three types as TRP candidates that are described as follows: a. peptides contain K residues that carry dimethylation or methylation and are not located at the C-terminus; b. peptides contain K, and the K does not carry dimethylation or methylation modification and are located at the C-terminus; c. peptides may not contain K, but the amino acid next to the N-terminal residue is K. Accessibility of peptides that can fit into the above types can be probed by TRAP, and are thus considered for TRP screening.

#### Determination of TRPs for drugs/metabolites

For quantitative screening of the peptides that exhibit significant abundance changes between the control and ligands (drugs/metabolites)-treated groups, we first classified them into two categories as loose and compact. The loose category refers to the peptides that become more chemically accessible to TRAP labeling reagents CD_2_O after given drugs/metabolites incubation. The increased chemical accessibility is detected by increased abundance of peptides that contain dimethylated K (type a) or decreased abundance of peptides belonging to type b and c. Conversely, the compact category refers to the peptides that become less accessible to TRAP labeling after given drugs/metabolites incubation. The reduced chemical accessibility is judged by the decreased abundance of type a peptides or increased abundance of type b and c peptides.

We set the standard of TRP screening for metabolites as peptides that display significant abundance changes with q value < 0.03 (q values are used to adjust for multiple testing using the Benjamini-Hochberg method to control the FDR at the cut-off level of 0.03) and the TRAP ratio (treated/control) > 1.5 or < 0.67 for compact peptides and TRAP ratio > 2 or < 0.5 for loose peptides. We posit that loose peptides reflect indirect binding events due to primary target engagement, so more strict restriction was given for this category. For drug targets discovery made based on TMT-based multiplexed TRAP data, a more stringent criteria of TRPs was utilized and the screening standard of drug TRPs was set as peptides that display q value< 0.01 and TRAP ratio > 2 or <0.5 in the presence of ligands relative to a control for both compact and loose peptides. As for target screening based on label-free quantification data, the screening standard of TRPs for metabolites was set as peptides that display p value< 0.05 (by Student’s t-test) and TRAP ratio > 1.5 or < 0.67 in the presence of metabolites relative to control, while for drugs a criteria of p value < 0.01 and TRAP ratio > 2 or < 0.5 was used.

#### Quality assessment of BAPP results

In order to assess the quality of our results, we estimated true positive rate generated by the TRAP approach by modifying previous assessment method^8^. We collected the known interactions from the BRENDA repository (http://www.brenda-enzymes.org/) and set the species as *homo sapiens*. Among all potential enzyme-metabolite interactions, 140 known enzyme-metabolite interactions for *homo sapiens* in BRENDA were retrieved. Our TRAP results detected 30 known enzyme-metabolite interactions that are classified as true positive hits, which confer a true positive rate as 21.43% (calculated by 30/140).

#### Volcano plot of the TRAP-identified targetome

In the volcano plot, each point accords to a protein that is represented by a peptide selected based on a scoring system. The score of each quantified peptide for given proteins is obtained by consideration of both the TRAP ratio (treated/control) and p/q value of the peptide abundances between samples with and without given ligands shown as follows.

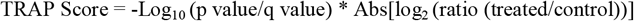

After scoring, the peptide with the maximum TRAP score was selected to represent the given protein.

#### Secondary structure analysis

UniProt identifiers of all quantified proteins were matched with PDB accession number from Protein Data Bank (PDB) (http://www.rcsb.org/pdb/search/searchModels.do) and only entries with >90% sequence identity were retrieved for analysis. Secondary structure information of the compiled protein pool was downloaded from the DSSP database (https://swift.cmbi.umcn.nl/gv/dssp) and Python script was used to extract the secondary structure for each quantified lysine residue. The extracted secondary structure classes were classified into four categories, namely helix (DSSP classes H, G, I), sheet (DSSP classes B, E), loop (T, S) and no structure (“”)^44^.

#### Conservation analysis

To estimate the sequence conservation of the obtained Lysine sites from TRPs, we calculated the Lysine sequence identity across 11 representative vertebrate species (Human, Rhesus monkey, Mouse, Rat, Cow, Dog, Opossum, Chicken, Frog, Zebrafish, and Fugu) using an in-house perl script. Specifically, we downloaded multiple amino acid sequence alignment of coding sequence (CDS) region across 100 species (multiz100way) from UCSC Genome Browser (Haeussler, et al. 2019). For each gene, its CDS region alignment across the selected 11 representative vertebrate species was further extracted. For each lysine from TRPs, its sequence conservation was estimated using the percentage of sequence conservation across 11 representative vertebrate species. To further examine whether the obtained TRP lysine sites were more conserved than random expectation, we calculated the sequence conservation of all quantified Lysine sites to estimate the background Lysine conservation, and then use Kolmogorov-Smirnov (KS) test to evaluate the statistical significance of excessive sequence conservation of the obtained TRP Lysine residues. The KS test p <0.05 was considered as significant.

#### Measurements of Euclidean distances

For functional sites analysis, PyMOL-Python scripts was used to measure the Euclidean distances between the atoms of lysine in TRPs and any atoms of annotated ligands (such as substrates, cofactors, products) in Å for enzymes assigned as glycolytic metabolites targets by TRAP for those with available structures retrieved from PDB files. The minimum distance was recorded to represent each ligand-target pair. Further, if the minimum distance is less than 10 Å, the lysine is categorized as functional and otherwise as unfunctional.

The active site boundary detectable by TRAP is defined based on the median of the minimum distances measured between the TRPs of known enzymatic targets that use the examined metabolites as substrates and their corresponding active sites.

For evaluation of the metabolites’ influence on given enzymes’ activities, the minimum distance between all TRPs from the identified targets involved in the carbohydrate metabolism pathway and their corresponding active sites were measured and shown in **Fig. 2k**.

#### GO and KEGG pathway analysis

All the TRAP-identified target proteins were annotated to non-overlapping GO molecule function (MF) terms. The MF classification was performed using the functional annotation tool of PANTHER (http://pantherdb.org/). GO MF terms include Transcription regulator activity (GO: 0140110), Catalytic activity (GO:0003824), Transporter activity (GO:0005215), Molecular transducer activity (GO:0060089), Translation regulator activity (GO:0045182), Structural molecule activity (GO:0005198), Molecular function regulator (GO:0098772), Binding (GO:0005488). If a protein belongs to more than one class, they are categorized based on the above order to prioritize. Proteins that do not match any of the above GO terms were sorted into the category “Uncategorized”.

Specifically, for proteins sorted in the “Catalytic activity” category, ClueGO app included in Cytoscape was employed to perform the KEGG pathway annotation network analysis. A setting of group p value <0.001 and inclusion of at least three genes in each group was used for filtering.

GO biological process (BP) analysis of the druggable glycolytic targetome was performed by BiNGO app included in Cytoscape with a setting of group p value <1e-15.

#### DrugBank analysis for ligandability classification

The DrugBank database (v. 5.1.3 released on 2019-04-02; group “all_ target_polypeptide_ids”) were downloaded and used to classify the glycolytic targetome into DrugBank (ligandable) and non-DrugBank (unligandable) proteins.

#### Protein-Protein interface analysis

To determine whether the identified glycolytic target interactions possibly affect protein-protein complexes, lysines of all quantified peptides or of TRPs were queried against the interfacial amino acids predicted by the PrePPI database (https://bhapp.c2b2.columbia.edu/PrePPI/) using a C-PPISP method. Lysines with their flanking ± 5 amino acids were all examined based on the corresponding protein sequences in the UniProt database.

#### Analysis of lysine PTM

The PTM information of lysines from all quantified proteins was collected and compiled with a Python script from the iPTMnet database (https://research.bioinformatics.udel.edu/iptmnet). Glycolytic metabolite-target interaction is considered to potentially affect the lysine-bearing PTM status if a given lysine in TRPs is reported to carry PTM.

### Glycolytic Metabolites target validation

#### siRNA Transfection

Scrambled small interfering RNA (siRNA) targeting TRIM28 (sc-38550), ENO1 (sc-37007), MTHFD1 (sc-61082), GSTP1 (sc-72091) and control siRNA (scrambled siRNA, sc-37007) were purchased from Santa Cruz Biotechnology (USA). For RNA interference, HCT116 cells were transfected at 24 hr after plating using lipofectamine™ RNAiMAX reagent (Invitrogen, Carlsbad, CA, USA) according to the manufacturer’s instructions. After 48 hr transfection, cells were collected to verify the silencing efficiency. For cell proliferation assay, at 24 hr post-transfection, cells were treated with G3P at concentrations of 0.5, 0.75 and 1 mM for 24 hr. For p53 expression measurement, cells were pretreated with 10 mM lactate for 4 hr at 24 hr post-transfection and then treated with 10 μM GNE140 or 0.1 μM doxorubicin (Dox) for another 20 hr.

#### Cell viability and proliferation assay

For cell viability assay, cells were plated in 96-well plates. After attachment for 24 hr, cells were treated with individual metabolites for another 48 hr. Then a CCK-8 assay was conducted using the cell counting kit (Yeasen, Shanghai, China, cat. no. 40203ES60) according to the manufacturer’s instructions. For determination of proliferation, cells were cultured in 5 individual 96-well plates and treated with the assayed glycolytic metabolites-supplemented medium containing 2% FBS. At indicated time points, cells were stained with Hoechst 33342 (Beyotime Biotechnology, cat. no. C1022) and cell number was counted using Lionheart FX Automated Live Cell Imager (BioTek, Winooski, VT, USA).

#### Colony formation assay

For colony formation assay, ~2000 HCT116 cells were seeded per well in 6-well plates and incubated with different concentrations of G3P or control solvents for 10 days. The colonies were then fixed with 4% paraformaldehyde for 20 min and stained with crystal violet solution (Beyotime Biotechnology, cat. no. C0121), rinsed with PBS, and imaged using Leica DMI 3000B light microscope (Leica, Wetzlar, Germany) in a blinded manner.

#### Western blotting

Cells were lysed in RIPA lysis buffer (Beyotime Biotechnology, cat. no. P0013B) supplemented with protease inhibitor cocktail. The protein concentrations were then determined by the BCA assay. The lysates were diluted by 4× XT Sample Buffer (Bio-Rad, Hercules, USA. cat. no. 1610791), heated to 95 °C for 5 min, cooled and separated by 8%-12% sodium dodecyl sulfate polyacrylamide gel electrophoresis (SDS-PAGE). After being transferred onto polyvinylidene difluoride (PVDF) membranes (Bio-Rad, cat. no. 1620177), proteins were blocked using 5% non-fat dry milk in Tris-buffered saline with 0.1% Tween 20 detergent (TBST) and incubated with primary antibodies at 4 °C overnight. After being washed five times with TBST, the membranes were incubated with HRP-conjugated secondary antibodies for 1 hr at 37 °C. The immunoblotted bands were detected by the addition of horseradish peroxidase (HRP) substrate (Bio-Rad, cat. no. 170-5601) and captured on a ChemDoc XRS+ System (Bio-Rad). The primary antibodies used in this study include antibodies against p53 (Abcam, Cambridge, UK, cat. no. ab1101), β-tubulin (Bioworld Technology, MN, USA, cat. no. AP0064,), TRIM28 (Cell Signaling Technology, Danvers, MA, USA, cat. no. 4123), cleaved PARP (Cell Signaling Technology, cat. no. 5625), cleaved caspase-3 (Cell Signaling Technology, cat. no. 9661), ENO1 (Cell Signaling Technology, cat. no. 3810), GSTP1 (Proteintech, Wuhan, China, cat. no. 15902-1-AP) and MTHFD1 (Proteintech, cat. no. 10794-1-AP).

#### Immunoblotting of proteome pan-acetylation

HCT 116 cells were plated at a density of 1.5 ×10^6^ cells/petri dish and incubated overnight at 37 °C with 5% CO_2_. After 24 hr, cells were cultured with given metabolites for 4 hr. The deacetylase inhibitor trichostatin A (TSA) at 0.5 μM or control solvent was added to the cultured medium for 24 hr, respectively. Cells were washed in PBS, suspended in lysis buffer containing 50 mM Tris, 150 mM NaCl, 1% Nonidet P-40 (NP-40) and protease inhibitors cocktail (pH 7.4). The lysates were incubated on ice for 30 min and sonicated with three 5-sec bursts. Thereafter, the lysates were centrifuged at 18,000 g at 4☐°C for 10 min, and the resultant supernatants were collected. A 50 μg aliquot of whole proteins was resolved on 10 % or 15 % SDS-PAGE gel according to the described Western blotting methods except 5% bovine serum albumin was used for blocking. A pan-acetylation antibody (Cell Signaling Technology, cat. no. 9441) was used as the primary antibody and equal loading amount of proteins for each sample was verified by Coomassie blue staining (Beyotime Biotechnology, cat. no. P0017).

#### Subcloning and mutagenesis

The synthetic genes encoding wild type (WT) human ENO1 (UniProt: P06733) and PKM2 (UniProt: P14618) with an N-terminal hexahistidine purification (His6) tag were subcloned into the GV296 vector (Genechem, Shanghai, China). The synthetic genes encoding WT human TRIM28 with an N-terminal maltose binding protein (MBP) affinity tag (UniProt: Q13263) were subcloned to the pMAL-p2X vector (New England Biolabs, Beverly, MA, USA).

The ENO1 K330E and TRIM28 K337E/K340E double mutants were obtained by site-directed mutagenesis. PCR primer sequences used for ENO1 and TRIM28 mutagenesis are listed as below:

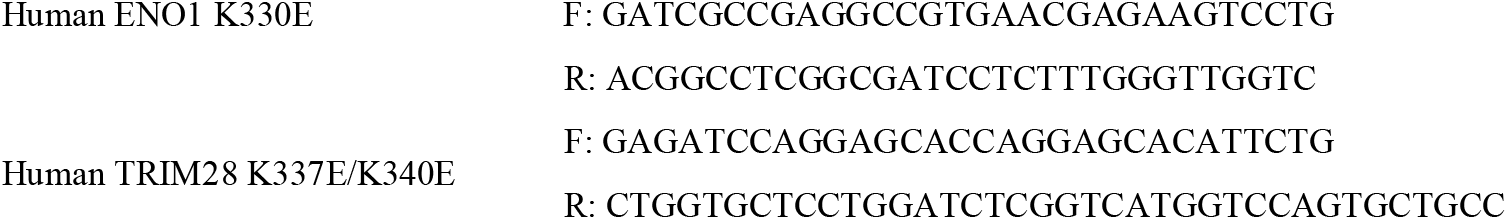

#### Recombinant protein expression and purification

All His-tagged recombinant protein expression and purification steps are generally as follows: plasmids were transformed into BL21(DE3) *E. coli* cells and the transformed bacteria were selected on LB plate containing 50 μg/mL kanamycin. The isolated colonies were grown in LB medium at 37 °C and 220 rpm until the OD_600_ of culture arrive 0.8. The target protein expression was induced with 0.5 mM isopropyl β-D-thiogalactoside (IPTG) by shaking overnight at 16 °C. Cells were harvested by centrifugation followed by resuspension in ultrapure water containing protease inhibitor cocktail, and then lysed by sonication. Lysates were centrifuged for 20 min at 18,000 g at 4 °C and the resultant supernatants were discarded. The His-tagged target proteins were then purified with a His-tag Protein Purification Kit (Beyotime Biotechnology, cat. no. P2226). After purification, proteins were concentrated by filter with a 10 kDa cut-off and the purity was evaluated by SDS-PAGE.

For the expression of TRIM28 constructs, cultures were supplemented with LB medium containing 50 μg/mL ampicillin and 0.5% glucose. Protein expression was induced at OD_600_ = 0.8 with 0.2 mM IPTG by shaking at 100 rpm overnight at 16 °C. To purify the MBP-tagged TRIM28, cells were resuspended in lysis buffer (50 mM Tris pH 7.4, 50mM NaCl, 5 mM DTT, 1:10,000 (v/v) benzonase solution, 1×protease inhibitor cocktail and lysed by sonication. The lysate was clarified by centrifugation. The supernatant was transferred to amylose resin (New England Biolabs) and incubated overnight at 4 °C. The protein was purified using a New England Biolabs kit according to the manufacturers’ protocol. The bound fusion protein was eluted with 10 mM maltose (New England Biolabs). The yield of the fusion proteins was evaluated by separation on SDS-PAGE and visualization via Coomassie blue staining.

#### PKM2 activity assay

*In vitro* activity of PKM2 was performed using luminescent Kinase-Glo Plus reagent (Promega, cat. no. V3773,) and used according to the manufacturer’s instructions. In brief, human recombinant PKM2 was diluted in the assay buffer (50 mM Tris HCl, 100 mM KCl, 10 mM MgCl_2_, PH 7.4) and incubated with each metabolite/drug/vehicle at 25 °C for 40 min. Substrate solution was prepared by mixing ADP at 200 μM and PEP at 200 μM with the assay buffer. Then, a 50 μL aliquot of substrate solution was added to 50 μL recombinant PKM2 solution, and allowed to react for 10 min at room temperature in 96-well plates. The luminescence response of each sample was measured due to the formation of ATP at 37 °C in a Gen5 platform (BioTek). The measurements were fitted by a nonlinear fitting algorithm (log [agonist] *vs.* response-variable slope with 4 parameters) in Prism 8.0.1 (GraphPad, San Diego, CA, USA).

#### GAPDH activity assay

*In vitro* GAPDH enzymatic activity assay was performed in 96-well plates at room temperature by measuring the reduced NAD^+^ level. The recombinant GAPDH (Abnova, Taiwan, China, cat. no. P4547) was first diluted in 10 mM sodium pyrophosphate buffer (pH 8.5) to 30 U/mg. A 100 μL of reaction mixture containing 20 mM sodium arsenate, 1 mM NAD^+^ and 2.88 mM G3P was readily added to GAPDH. The NAD^+^ level was examined by measuring the absorbance at 340 nm every 20 s for 20 min. Reaction rate was calculated by curve fitting of the time course measurements by linear regression.

#### ENO1 activity assay

*In vitro* ENO1 enzymatic activity is similar to PKM2 activity assay except that the substrate was changed to a substrate-enzyme mixture consisting of 400 μM 2PG, 400 μM ADP and 0.12 mg/mL recombinant PKM1. IC_50_ of G3P was determined in GraphPad Prism by a nonlinear fitting algorithm (log [inhibitor] *vs.* response-variable slope with 4 parameters).

#### NAMPT activity assay

*In vitro* activity of NAMPT was measured using a NAMPT colorimetric assay kit (CycLex, MBL International, cat. no. CY 1251V2) in 96-well plates by the one-step method. In brief, recombinant NAMPT was diluted in NAMPT assay buffer and subsequently incubated with solvent/metabolites/FK866 for 30 min at 30 °C. The reaction was initiated by the addition of 60 μL One-Step Assay Mixture to each well. NAMPT activity was measured photometrically in absorbance at 450 nm for 60 min with a 1 min-interval.

#### Thermal shift assay and isothermal dose-response assay

For thermal shift assay conducted using cell lysate samples, cultured HCT116 cells were harvested and washed with PBS. The washed cells were diluted in 0.1% NP-40 lysis buffer supplemented with 1× protease inhibitor cocktail and 1× Phosphatase Inhibitor Cocktail. The cell suspensions were freeze-thawed five times using liquid nitrogen and passed through a 27” gauge needle five times. The samples were snap-frozen in liquid nitrogen for > 1 min and placed onto a heating block set at 25 °C until 60% of the confluent was thawed. Then, the samples were subjected to centrifugation at 20,000 g for 20 min at 4 °C. The supernatant was collected and divided into two aliquots, with one aliquot being treated with given glycolytic metabolites and another aliquot with the solvent as the control. After 60 min-incubation at room temperature, each sample were further divided into 8 aliquots and heated individually at their designated temperature for 3 min in a 96-well thermal cycler followed by cooling at room temperature for 3 min. The heated lysates were centrifuged at 20,000 g for 20 min at 4 °C in order to separate the soluble fractions from precipitates. Each resultant soluble fraction of the proteome was transferred to a new low-adsorption 1.5 mL microtube and analyzed by SDS-PAGE followed by western blotting for target proteins.

For the cell lysate-level isothermal dose-response (ITDR) experiments, G3P were serially diluted to generate a 9 points dose–response curve. HCT116 cells lysate were incubated with the assayed metabolite of serial concentrations (at least eight concentrations) and one vehicle as control in 1.5 mL low absorption microtubes for 1 hr at room temperature. Then, the samples were aliquoted to 120 μL and transferred into 200 μL PCR microtubes followed by heating at designated temperatures for 3 min in a 96-well thermal cycler and subsequent cooling for 3 min at room temperature. Then, isolation of the soluble fractions and immunoblotting of given target protein are repeated as described for thermal shift assays.

For cell lysate-level thermal shift assay, the band intensities detected at increasing temperatures were normalized to that of the lowest temperature, and the Boltzmann sigmoid equation was fitted using GraphPad Prism. For the ITDR experiments, the band intensities at increasing doses were normalized to the intensity at the highest concentration of the assayed metabolites and analyzed by the saturation binding curve function in Prism.

#### Surface plasmon resonance analysis

SPR analysis was conducted on a Biacore T200 system (GE Healthcare, Sweden). Target protein was diluted in 10 mM sodium acetate and immobilized via the amine coupling method on a CM5 sensor chip. Metabolites were dissolved in H_2_O and diluted to a serial concentration with running buffer (PBS with 0.05% tween 20). Then, the metabolites were injected through the reference and active channels at a flow rate of 30 μL/min. The association and dissociation times were both set at 60 s. The affinity fitting was carried out on a Biacore T200 evaluation software by global fitting via a steady-state affinity model to obtain the equilibrium dissociation constant K_D_.

#### Protein Thermal stability screen by nanoDSF

Both the WT-ENO1 (0.5 μg/μL) and the mutant-ENO1 (0.5 μg/μL) were incubated with 2 mM G3P for 30 minutes firstly. Samples were filled within the nanoDSF capillaries (n=3), and subsequently loaded into the Prometheus NT.48 device (NanoTemper Technologies, Germany). Samples were heated from 20 °C to 95 °C with a slope of 1 °C/min and the unfolding transition temperatures were automatically identified by the PR. ThermControl software (NanoTemper Technologies). Raw data was exported for plotting the thermal stability curve in GraphPad Prism.

#### Relative quantification of metabolites by multiple reaction monitoring analysis

HCT 116 cells were plated onto 6-well plates. After attachment for 24 h, cells were incubated with NaF (2mM) and GNE140 (10 μM) for another 24 hr in serum-free medium. Metabolites were extracted with 1 mL pre-cooled methanol-water (8:2 by volume) containing chloro-phenylalanine as the internal standard. Samples were dried, resuspended in H_2_O and then measured by MRM analysis.

Quantitative analysis of the metabolites was performed on a QTRAP 5500 mass spectrometer (SCIEX) equipped with a Shimadzu LC-30A HPLC system. The extracted metabolites were separated on an XBridge BEH Amide HPLC column (100 mm × 4.6 mm, 3.5 μm) (Waters). The mobile phase consisted of solvent A (95% 5 mM ammounium acetate buffer, pH adjusted to 9, 5% acetonitrile) and solvent B (ACN). The gradient was set as follows: 0-3 min, 85% B; 3-6 min, 85-30% B; 6-15 min, 30-2% B; 15-18 min, 2%B; 18-19 min, 2-85% B; 19-26 min, 85% B. The flow rate was set at 0.4 mL/min. The ESI source conditions on the 5500 QTRAP system were set as follows: curtain gas 40☐psi, nebulizer gas 25☐psi, IonSpray voltage −4500☐V, ion source temperature 600 °C and CAD gas medium. The multiple reaction monitoring (MRM) transitions parameters for quantitative analysis of the glycolytic metabolites were modified based on previously reported declustering potential (DP), entrance potential (EP) and collision energy (CE)^45^. Data processing was carried out using Analyst software (SCIEX, version 1.6.1).

#### Flow cytometric analysis of cell apoptosis

Cell apoptotic rate was examined using an Annexin-V/FITC Kit (BD Biosciences, San Jose, CA, USA). Cells were pretreated with pyruvate for 4 hr and then treated with TSA for 24 hr. The cells were harvested, washed with PBS, resuspended in 1× Annexin V binding buffer, and stained with annexin V and PI for 15☐min at room temperature in darkness. The rates of apoptotic cells were determined using an Accuri C6 flow cytometer (BD Biosciences).

#### Homology Modeling

The 3D structure of human MTHFD1 (residues 301-935) was modeled according to the crystal structure of formate-tetrahydrofolate ligase (*moorella thermoacetica*, PDB code: 4IOJ1), which shows the sequence identity of 48.56% (and similarity of 61.7%) to the human MTHFD1 and represents the most similar crystal structure to human MTHFD1 in Protein Data Bank (2020.02.09). Modeller9.232 was used for the structure construction, and amber ff14SB force field3 was employed for the structure refinement, in which all the heavy atoms in the protein backbone were constrained with 5 kcal/mol·Å2 with other atoms free for moving. All the structure refinement was conducted with pmemd in AmberTools18.5.

#### Statistical analysis

Statistical analysis was performed in GraphPad Prism. All data represent mean ± SEM (n = 3 or 5 per group as indicated in legend). The statistical significance of differences between two groups was determined using Student’s t-test (unpaired, two-tailed) unless otherwise specified, *p < 0.05, **p < 0.01, ***p < 0.001, ****p < 0.0001.

## DATA AND SOFTWARE AVAILABILITY

The mass spectrometry proteomics data have been deposited to the ProteomeXchange Consortium (http://proteomecentral.proteomexchange.org) via the iProX partner repository^46^ with the dataset identifier IPX0002602000/PXD022568.

## REFERENCES

1. Fu, X. et al. 2-Hydroxyglutarate inhibits ATP synthase and mTOR signaling. Cell Metab. 22, 508–515 (2015).

2. Tang, W. H. et al. Intestinal microbial metabolism of phosphatidylcholine and cardiovascular risk. N Engl J Med. 368, 1575–1584 (2013).

3. Platten, M. et al. Tryptophan metabolism as a common therapeutic target in cancer, neurodegeneration and beyond. Nat Rev Drug Discov. 18, 379–401 (2019).

4. Zaslona, Z. et al. Cytokine-like roles for metabolites in immunity. Mol Cell. 78, 814–823 (2020).

5. Zhang, D. et al. Metabolic regulation of gene expression by histone lactylation. Nature. 574, 575–580 (2019).

6. Zhang, W. et al. Lactate is a natural suppressor of RLR signaling by targeting MAVS. Cell. 178, 176–189 (2019).

7. Mills, E. L. et al. Itaconate is an anti-inflammatory metabolite that activates Nrf2 via alkylation of KEAP1. Nature. 556, 113–117 (2018).

8. Piazza, I. et al. A map of protein-metabolite interactions reveals principles of chemical communication. Cell. 172, 358–372 (2018).

9. Niphakis, M. J. et al. A global map of lipid-binding proteins and their ligandability in cells. Cell. 161, 1668–1680 (2015).

10. Hulce, J. J. et al. Proteome-wide mapping of cholesterol-interacting proteins in mammalian cells. Nat Methods. 10, 259–264 (2013).

11. Bollong, M. J. et al. A metabolite-derived protein modification integrates glycolysis with KEAP1-NRF2 signalling. Nature. 562, 600–604 (2018).

12. Qin, W. et al. S-glycosylation-based cysteine profiling reveals regulation of glycolysis by itaconate. Nat Chem Biol. 15, 983–991 (2019).

13. Sridharan, S. et al. Proteome-wide solubility and thermal stability profiling reveals distinct regulatory roles for atp. Nat Commun. 10, 1155 (2019).

14. Schulze, A. et al. How cancer metabolism is tuned for proliferation and vulnerable to disruption. Nature. 491, 364–373 (2012).

15. Moellering, R. E. et al. Functional lysine modification by an intrinsically reactive primary glycolytic metabolite. Science. 341, 549–553 (2013).

16. Lomenick, B. et al. Target identification using drug affinity responsive target stability (DARTS). Curr Protoc Chem Biol. 3, 163–180 (2011).

17. Huber, K. V. et al. Proteome-wide drug and metabolite interaction mapping by thermal-stability profiling. Nat Methods. 12, 1055–1057 (2015).

18. Reinhard, F. B. et al. Thermal proteome profiling monitors ligand interactions with cellular membrane proteins. Nat Methods. 12, 1129–1131 (2015).

19. Molina, D. M. et al. Monitoring drug target engagement in cells and tissues using the cellular thermal shift assay. Science. 341, 84–87 (2013).

20. West, G. M. et al. Quantitative proteomics approach for identifying protein-drug interactions in complex mixtures using protein stability measurements. Proc Natl Acad Sci USA. 107, 9078–9082 (2010).

21. Raines, R. T. Ribonuclease A. Chem Rev. 98, 1045–1066 (1998).

22. Dombrauckas, J. D. et al. Structural basis for tumor pyruvate kinase M2 allosteric regulation and catalysis. Biochemistry. 44, 9417–9429 (2005).

23. Anastasiou, D. et al. Pyruvate kinase M2 activators promote tetramer formation and suppress tumorigenesis. Nat Chem Biol. 8, 1008–1008 (2012).

24. Bamberger, T. C. et al. Covalent protein painting reveals structural changes in the proteome in alzheimer disease. bioRxiv. (2020). doi: https://doi.org/10.1101/2020.01.31.929117.

25. Kornberg, M. D. et al. Dimethyl fumarate targets GAPDH and aerobic glycolysis to modulate immunity. Science. 360, 449–453 (2018).

26. Lambert, S. A. et al. The human transcription factors. Cell. 172, 650–665 (2018).

27. Czerwinska, P. et al. The complexity of TRIM28 contribution to cancer. J Biomed Sci. 24, 63 (2017).

28. Jurchott, K. et al. Identification of Y-box binding protein 1 as a core regulator of MEK/ERK pathway-dependent gene signatures in colorectal cancer cells. PLoS Genet. 6 (2010).

29. Qin, G. et al. NPM1 upregulates the transcription of PD-L1 and suppresses T cell activity in triple-negative breast cancer. Nat Commun. 11, 1669 (2020).

30. Zhang, J. et al. Histone deacetylase inhibitors and cell death. Cell Mol Life Sci. 71, 3885–3901 (2014).

31. Liau, N. P. D. et al. Negative regulation of RAF kinase activity by ATP is overcome by 14-3-3-induced dimerization. Nat Struct Mol Biol. 27, 134–141 (2020).

32. Molzan, M. et al. Synergistic binding of the phosphorylated S233- and S259-binding sites of C-RAF to one 14-3-3zeta dimer. J Mol Biol. 423, 486–495 (2012).

33. Ganapathy K. S. et al. Tumor glycolysis as a target for cancer therapy: progress and prospects. Mol Cancer. 12, 152 (2013).

34. Ying, J. et al. The effect of sodium fluoride on cell apoptosis and the mechanism of human lung BEAS-2B cells in vitro. Biol Trace Elem Res. 179, 59–69 (2017).

35. Boudreau, A. et al. Metabolic plasticity underpins innate and acquired resistance to LDHA inhibition. Nat Chem Biol. 12, 779–786 (2016).

36. Christensen, K. E. et al. The MTHFD1 p.Arg653Gln variant alters enzyme function and increases risk for congenital heart defects. Hum Mutat. 30, 212–220 (2009).

37. Coles, B. F. et al. Detoxification of electrophilic compounds by glutathione S-transferase catalysis: determinants of individual response to chemical carcinogens and chemotherapeutic drugs? Biofactors. 17, 115–130 (2003).

38. Khan, J. A. et al. Molecular basis for the inhibition of human NMPRTase, a novel target for anticancer agents. Nat Struct Mol Biol. 13, 582–588 (2006).

39. Kastenhuber, E. R. et al. Putting p53 in context. Cell. 170, 1062–1078 (2017).

40. Tran, T. Q. et al. Α-ketoglutarate attenuates wnt signaling and drives differentiation in colorectal cancer. Nat Cancer. 1, 345–358 (2020).

41. Leone, R. D. et al. Glutamine blockade induces divergent metabolic programs to overcome tumor immune evasion. Science. 366, 1013–1021 (2019).

## REFERENCES

42. Chen, W. W. et al. Absolute quantification of matrix metabolites reveals the dynamics of mitochondrial metabolism. Cell. 166, 1324–1337 (2016).

43. Miyo, M. et al. Metabolic adaptation to nutritional stress in human colorectal cancer. Sci Rep. 6, 38415 (2016).

44. Feng, Y. H. et al. Global analysis of protein structural changes in complex proteomes. Nat Biotechnol. 32, 1036–1044 (2014).

45. Yuan, M. et al. A positive/negative ion-switching, targeted mass spectrometry-based metabolomics platform for bodily fluids, cells, and fresh and fixed tissue. Nat Protoc. 7, 872–881 (2012).

46. Ma, J. et al. iProX: an integrated proteome resource. Nucleic Acids Res. 47, 1211–1217 (2019).

