## Extended Data for "Chemoproteomics Maps Glycolytic Targetome in Cancer Cells" (from another version of this preprint: version 1, 2020-11-19)

### Figure S1

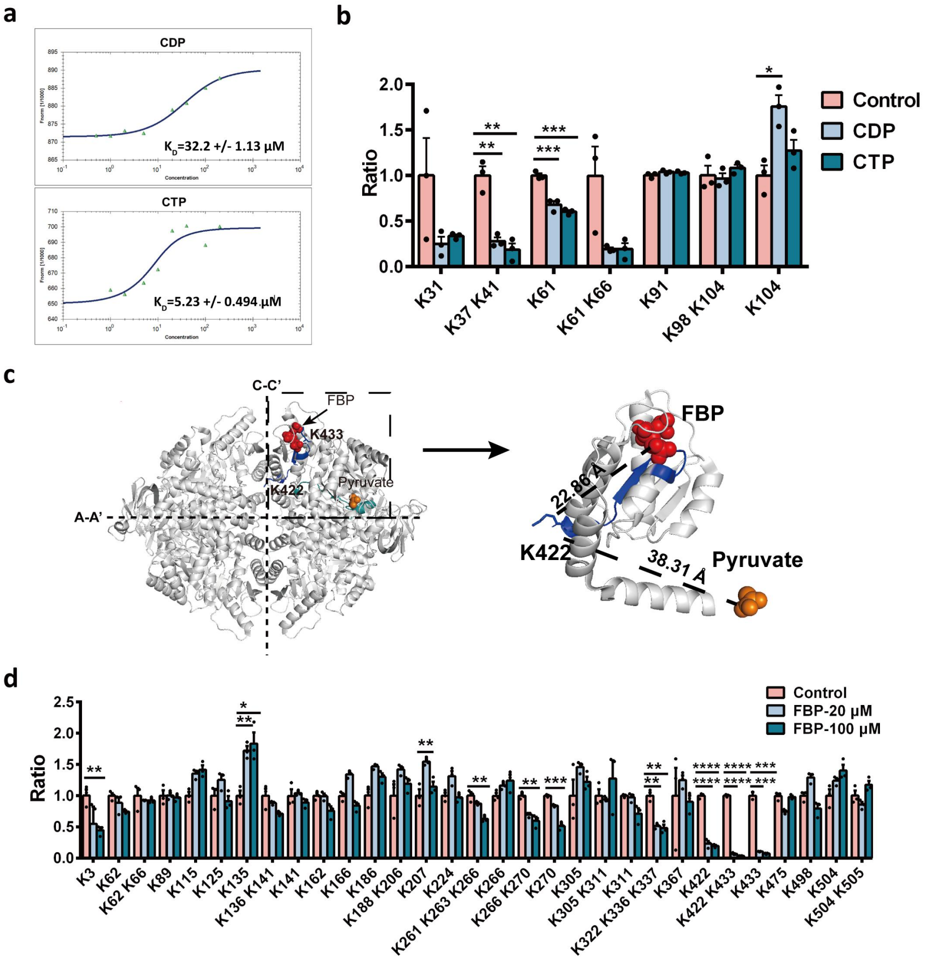


**Figure S1. Benchmarking the TRAP approach in probing ligand-target interactions with model proteins including the purified RNase A and recombinant PKM2.**

**(a)** Microscale thermophoresis (MST) of RNase A shows its ligand CTP possessing stronger affinity compared to its ligand CDP.

**(b)** A summary of accessibility changes of quantified lysines in RNase A in response to CDP/CTP binding via the TRAP method.

**(c)** Tetramerization of PKM2 (grey, cartoon) induced by the endogenous glycolytic metabolite FBP (red, sphere) by allosteric mechanism (PDB: 4B2D). Detailed X-ray structure of PKM2 shows the $K^{422}$ residue (blue, stick) is distant from FBP and pyruvate (orange, sphere) based on the measured minimal Euclidean distances in Angstroms between $K^{422}$ and FBP/pyruvate (indicated in dashed lines in insert).

**(d)** A summary of accessibility changes of quantified lysines in the human recombinant PKM2 in response to FBP binding is obtained via the TRAP method.

Data represent mean ± SEM (n = 3), two-tailed t-test, *P < 0.05, **P < 0.01, ***P < 0.001, ****P < 0.0001.

### Figure S2

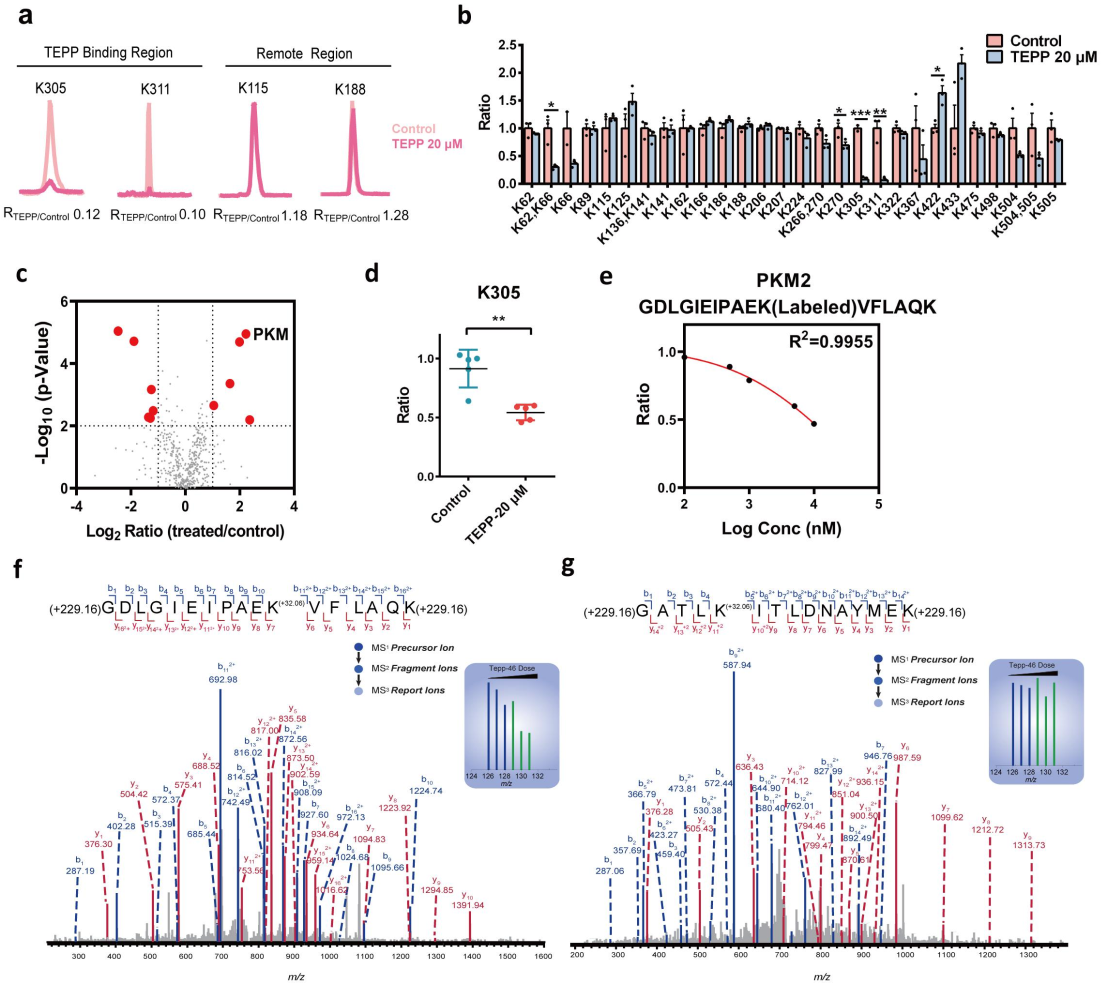


**Figure S2. TRAP identifies the TEPP-46 binding protein as PKM2 and pinpoints the binding site from samples including recombinant PKM2 and complex biological lysates.**

**(a)** TRAP identifies the proximal binding sites of TEPP-46, $K^{305}$ and $K^{311}$, from the recombinant PKM2 based on the significantly decreased peak abundances of TRPs following TEPP-46 incubation (n=3). The annotated TRAP Ratio is averaged from TRPs and non-TRPs examined from the three replicates, and the representative extracted ion chromatogram (EICs) of these peptides located in different domains are shown.

**(b)** A summary of accessibility changes of quantified lysines in human recombinant PKM2 in response to TEPP-46 binding via the TRAP method (n=3).

**(c)** Volcano plots identify the binding target PKM2 for TEPP-46 from *E. coli* lysates (n=5). Proteins displaying significantly altered accessibility change following TEPP-46 administration with the significance cutoff of p<0.01 and the TRAP ratio (treated/control) cutoff of >2 or <0.5 are highlighted in red.

**(d)** Relative abundance of the $K^{305}$-containing peptide is significantly decreased in *E. coli* lysate following TEPP-46 treatment (n=5).

**(e)** The TRAP ratio of the $K^{305}$-containing TRP indicates this region displaying dose-dependent decrease of accessibility when the incubated TEPP-46 concentration was increased. (fitted by dose-response-inhibition equation in Graphpad). Specifically, TMT 6-plex reagents were used to label the HCT116 cell lysates incubated with control solvent and TEPP-46 at serially increased concentrations (100 nM, 500 nM, 1 μM, 5 μM, 10 μM).

**(f)** Representative MS3-based TMT reporter ions showed that the $K^{305}$-containing TRP in PKM2 exhibits dose-dependent decreased accessibility in response to increased TEPP-46 concentration.

**(g)** Representative MS3-based TMT reporter ions showed that a non-TRP in PKM2 exhibits negligible changes in response to increased TEPP-46 concentration.

Data represent mean ± SEM (n = 3), two-tailed t-test, *P < 0.05, **P < 0.01, ***P < 0.001.

#### Figure S3

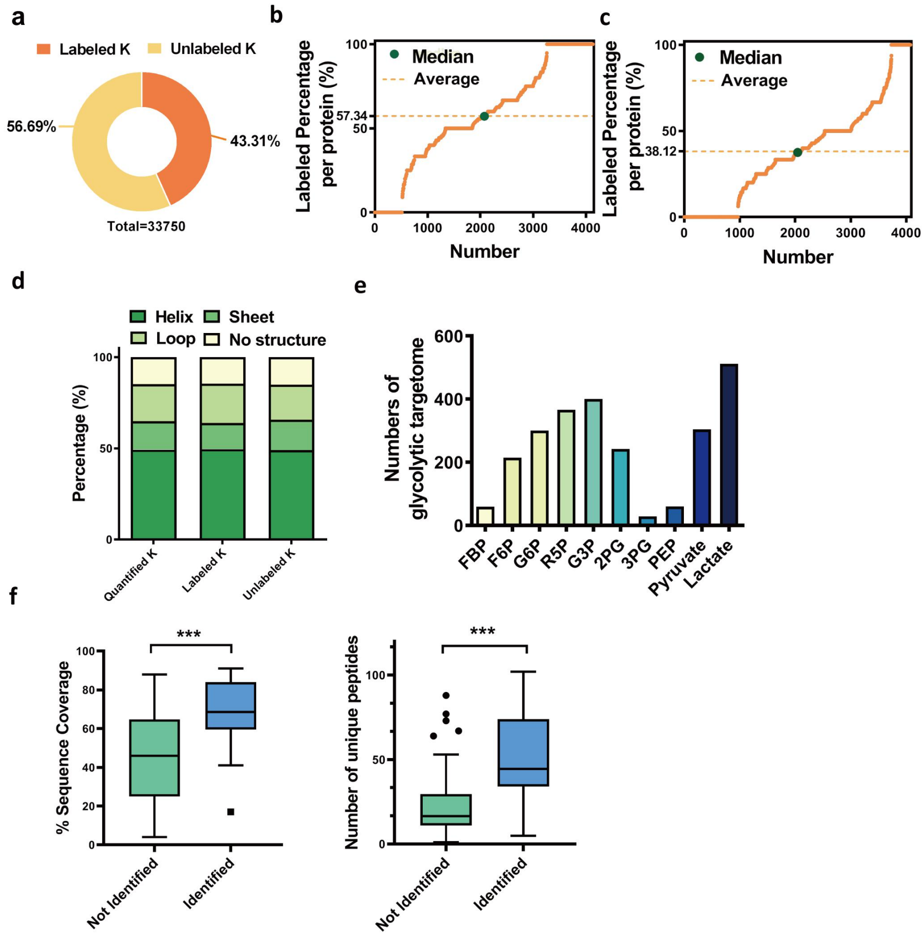


**Figure S3. Characterization of the overall multiplexed-TRAP data that assigned glycolytic targetome from the model cancer cell milieu.**

**(a)** Broad labeling coverage of reactive lysines achieved by the dimethylation labeling employed by TRAP. The second batch of the multiplexed-TRAP data was used for plotting.

**(b)** Chemical accessibility of proteinaceous lysines is examined by the labeled fraction of lysine residues for each quantified protein using the first batch of the multiplexed-TRAP data.

**(c)** Chemical accessibility of proteinaceous lysines is revealed by the labeled fraction of lysine residues for each quantified protein using the second batch of the multiplexed-TRAP data.

**(d)** Labeling preference analysis for high-order structures probed by TRAP based on the second batch of the multiplexed-TRAP data.

**(e)** Number of glycolytic metabolite-binding protein interactions detected by TRAP.

**(f)** The detection of the TRAP-identified glycolytic metabolites targetome is biased by mass spectral undersampling of the proteome based on the comparison of the sequence coverage and the number of detected unique peptides between known glycolytic targets (retrieved from BRENDA, species: human) that can (19 proteins) or cannot be identified (76 proteins) by TRAP (two-tailed t-test, ***P < 0.001).

#### Figure S4

a

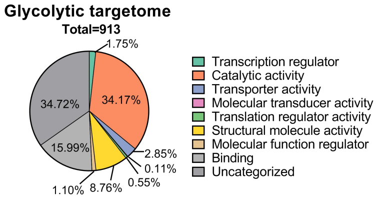


b

| | |
|---|---|
| **1. Metabolism** | |
| **1.1 Carbohydrate metabolism** | |
| **Glycolysis / Gluconeogenesis** | **KEGG:00010** |
| **Citrate cycle (TCA cycle)** | **KEGG:00020** |
| **Fructose and mannose metabolism** | **KEGG:00051** |
| **Pyruvate metabolism** | **KEGG:00620** |
| **Glyoxylate and dicarboxylate metabolism** | **KEGG:00630** |
| **1.5 Amino acid metabolism** | |
| **Cysteine and methionine metabolism** | **KEGG:00270** |
| **2. Genetic Information Processing** | |
| **2.1 Transcription** | |
| **Aminoacyl-tRNA biosynthesis** | **KEGG:00970** |
| **2.3 Folding, sorting and degradation** | |
| **Proteasome** | **KEGG:03050** |
| **2.4 Replication and repair** | |
| **DNA replication** | **KEGG:03030** |

c

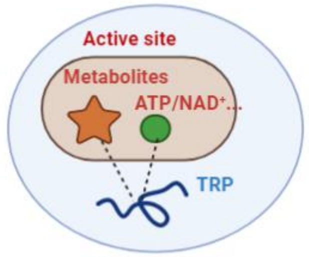


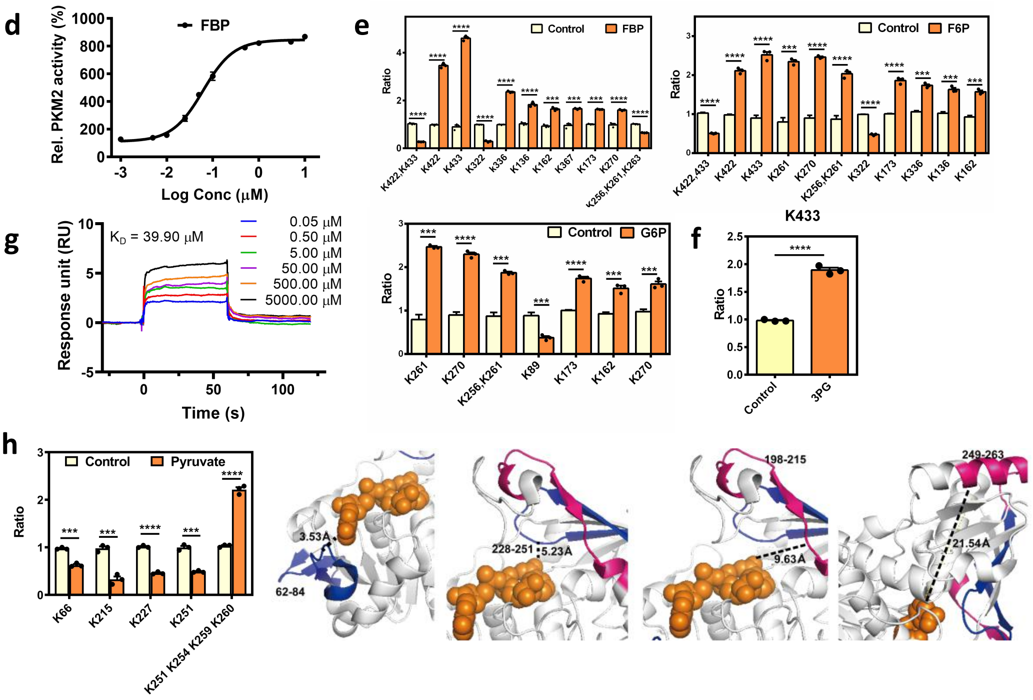


**Figure S4. Functional assessment and validation of the identified glycolytic metabolite-target interactions belonging to the carbohydrate metabolism pathway.**

**(a)** Gene Oncology (GO) Molecular Function (MF) analysis of the identified glycolytic targetome shows enrichment in proteins ascribed to catalytic activity.

**(b)** Enriched pathways of the glycolytic targetome ascribed to catalytic activity via the KEGG pathway annotation network analysis.

**(c)** Illustration of how the boundary of active site is defined via the multiplexed-TRAP approach. Briefly, the minimal Euclidean distance between the TRPs from the TRAP-identified targets that use the assayed metabolites as substrates and the corresponding active site (retrieved from PDB) were measured, and the resultant median of the collected distances is used to represent the active site boundary that can be probed by TRAP.

**(d)** Enzymatic activity of human recombinant PKM2 activated by FBP (n = 3).

**(e)** A summary of accessibility changes of the TRPs from PKM2 in HCT116 cells in response to the incubation with FBP, F6P and G6P, respectively (n=3)
**(f)** The TRAP ratio of the peptide containing $K^{433}$ of PKM2 in response to 3PG engagement (n=3).
**(g)** Surface Plasmon Resonance (SPR) assay determines the binding affinity of pyruvate to the human recombinant GAPDH.
**(h)** TRAP ratios (n=3) and the measured distance to the active site for all the pyruvate TRPs in GAPDH. Data represent mean ± SEM, two-tailed t-test, ***P < 0.001, ****P < 0.0001.

### Figure S5

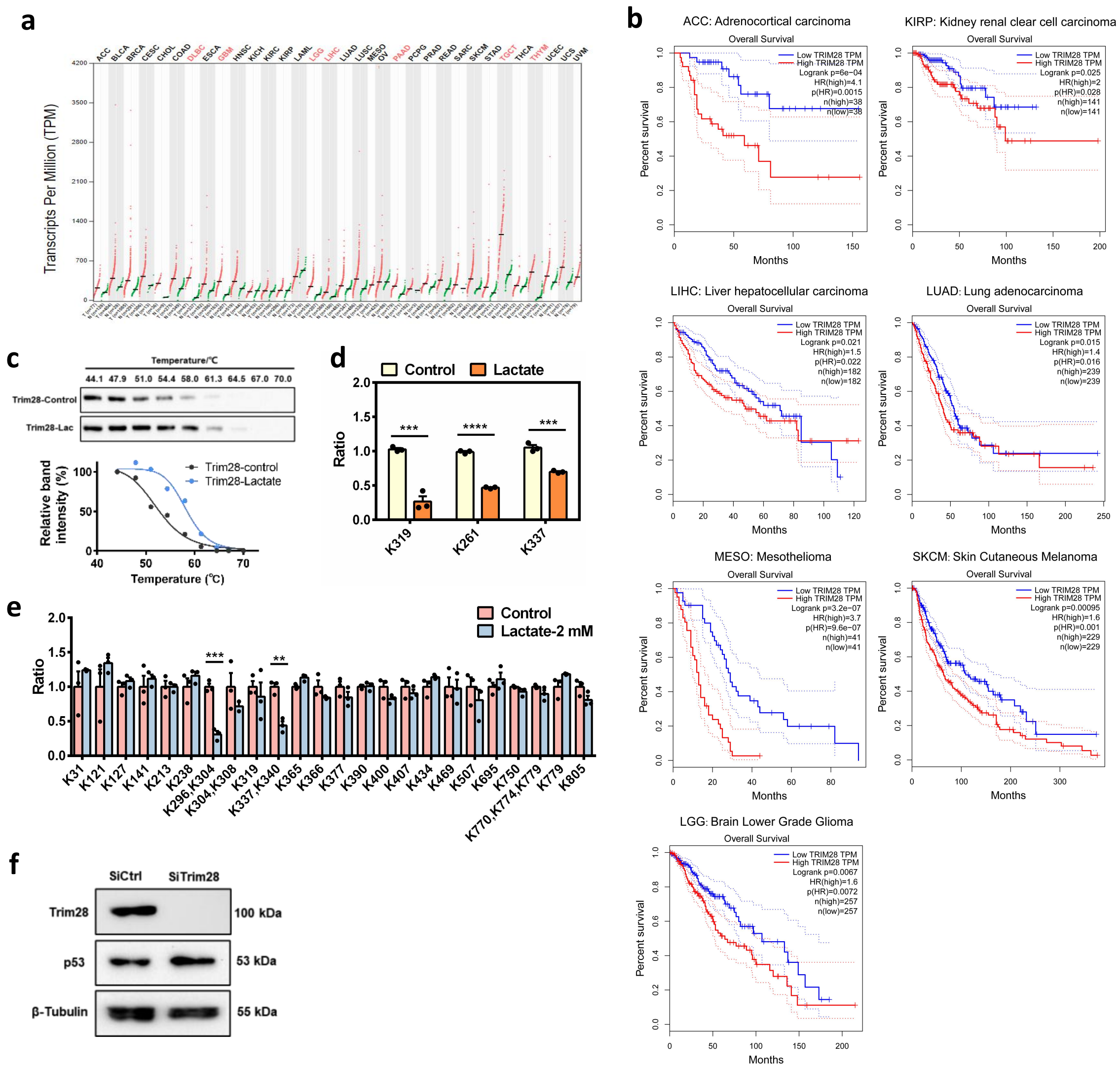


**Figure S5. Validation and characterization of the interaction between lactate and TRIM28.**

**(a)** Gene expression profile of TRIM28 across given cancer types and paired normal tissues plotted via GEPIA (http://gepia.cancer-pku.cn/detail.php?gene=trim28).

**(b)** High TRIM28 gene expression level is unfavored for patients' survival for the examined cancer types as summarized by GEPIA.

**(c)** Thermal shift assay validates the interaction between lactate and TRIM28 in HCT116 cell lysates..

**(d)** TRAP ratios of the lactate TRPs in TRIM28 obtained from the assayed HCT116 cell lysates. Data represent mean ± SEM (n = 3), two-tailed t-test, ***P < 0.001, ****P < 0.0001.

**(e)** Accessibility changes of the quantified lysines in human recombinant TRIM28 in response to lactate binding was probed via TRAP and summarized here. Data represent mean ± SEM (n = 3), two-tailed t-test, **P < 0.01, ***P < 0.001.

**(f)** Knockdown efficiency using siRNA targeting TRIM28 in HCT116 cells.

#### Figure S6

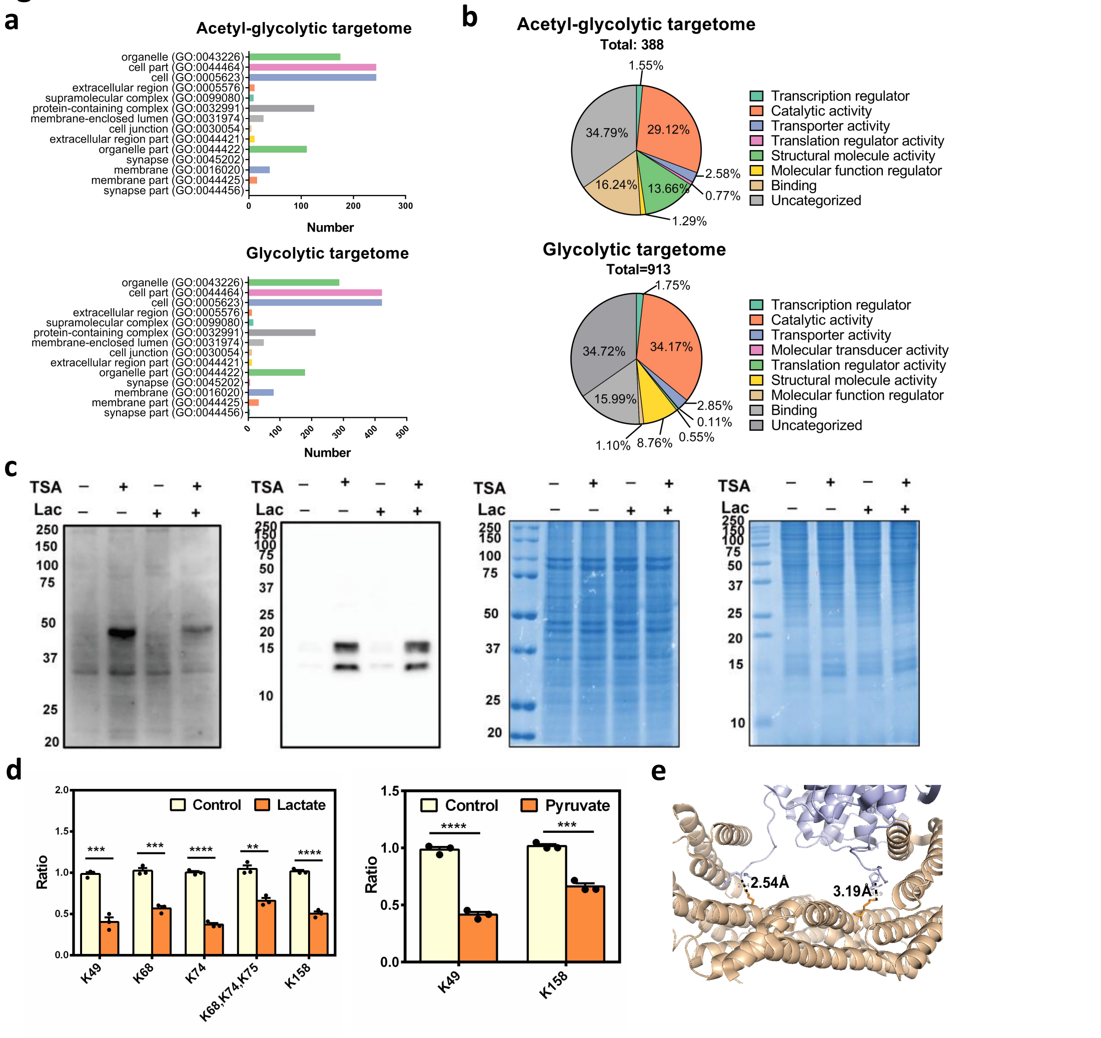


**Figure S6. Novel regulatory modalities of glycolytic metabolites involving protein acetylation and protein-protein interactions.**

**(a)** GO analysis shows diverse subcellular locations for the TRAP-assigned glycolytic targetome that have been documented by iPTMnet to carry acetylation. The subcellular distribution pattern resembles that of the whole glycolytic targetome.

**(b)** Distribution pattern of the molecular functions summarized for the whole glycolytic targetome and the target pool that have been reported to carry acetylation as retrieved from iPTMnet.

**(c)** Lactate administration (2 mM, 4 hr pretreatment) partially abrogated the protein pan-acetylation level upregulated by the co-administered trichostatin A (TSA, 0.5 μM, 20 hr) in HCT116 cells. The immunoblotted image and the Coomassie blue stained image of the relatively high molecular weight protein (left panels) and the relatively low molecular weight proteins (right panels) are shown.

**(d)** TRAP ratios of the pyruvate and lactate TRPs identified from the 14-3-3 protein zeta/delta in HCT116 cell lysates. Data represent mean ± SEM (n = 3), two-tailed t-test, $^{**}P < 0.01$, $^{***}P < 0.001$, $^{****}P < 0.0001$.

**(e)** Enlarged X-ray structure (PDB, 6U2H) of the 14-3-3 protein zeta/delta dimer (orange, cartoon) and B-RAF (light blue, cartoon) shows the minimal distances between the pyruvate/lactate-binding responsive site $K^{49}$ (wheat, stick) in the 14-3-3 protein zeta/delta dimer and B-RAF are 2.54 Å and 3.19 Å (measured along the black dashed line).

### Figure S7

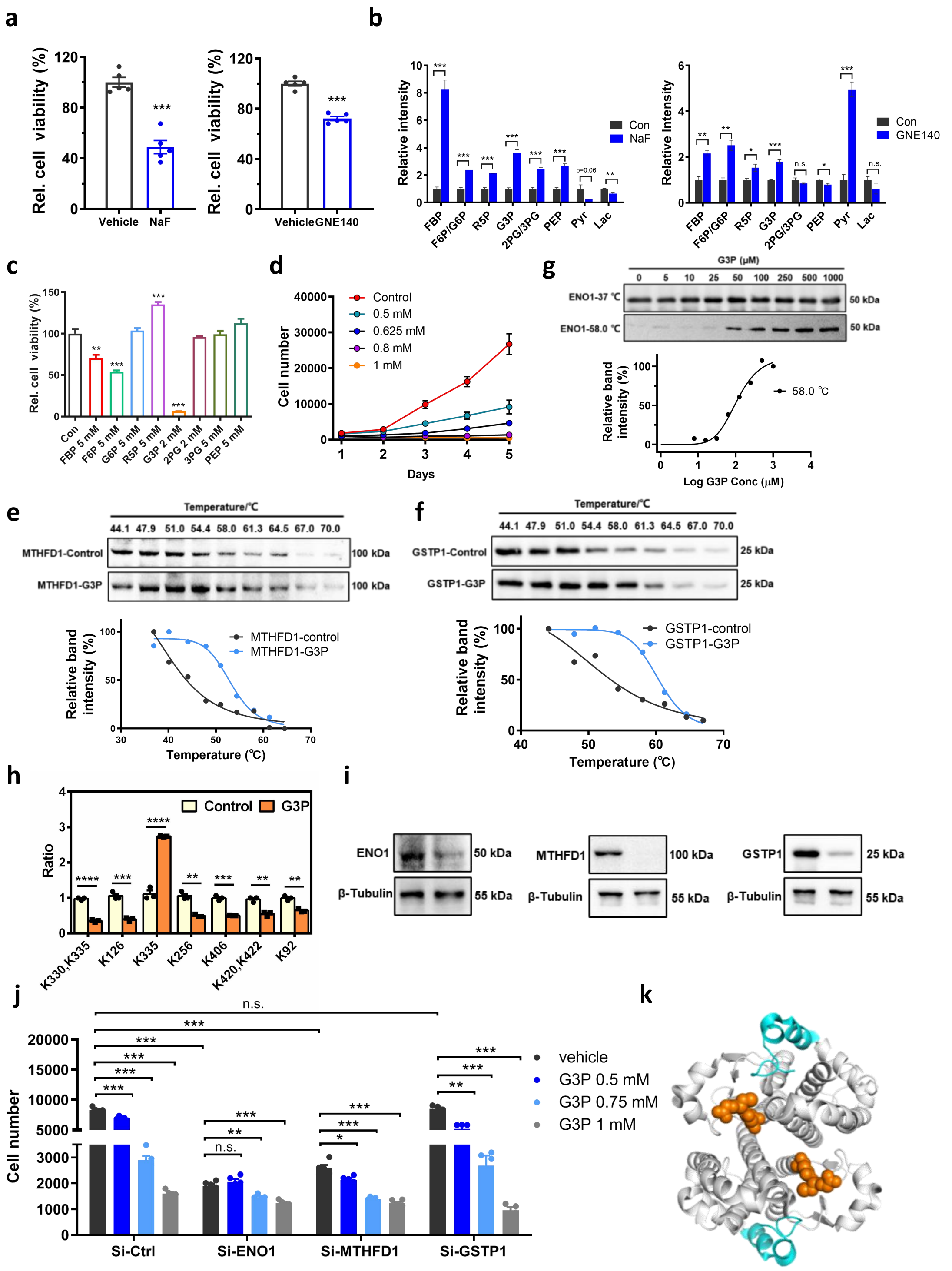
a
Rel. cell viability (%)
120
80
40
0
***
Vehicle
NaF
VehicleGNE140
b
Relative intensity
Con
NaF
GNE140
FBP
F6P/G6P
R5P
G3P
2PG/3PG
PEP
Pyr
Lac
p=0.06
n.s.
c
Con
FBP 5 mM
F6P 5 mM
G6P 5 mM
R5P 5 mM
G3P 2 mM
2PG 2 mM
3PG 5 mM
PEP 5 mM
d
Cell number
Days
Control
0.5 mM
0.625 mM
0.8 mM
1 mM
g
G3P (μM)
0
5
10
25
50
100
250
500
1000
ENO1-37 ℃
ENO1-58.0 ℃
50 kDa
Relative band intensity (%)
58.0 ℃
Log G3P Conc (μM)
e
Temperature/℃
44.1
47.9
51.0
54.4
58.0
61.3
64.5
67.0
70.0
MTHFD1-Control
MTHFD1-G3P
100 kDa
MTHFD1-control
MTHFD1-G3P
Temperature (℃)
f
GSTP1-Control
GSTP1-G3P
25 kDa
GSTP1-control
GSTP1-G3P
h
Ratio
Control
G3P
K330,K335
K126
K335
K256
K406
K420,K422
K92
i
ENO1
MTHFD1
GSTP1
β-Tubulin
55 kDa
j
Cell number
vehicle
G3P 0.5 mM
G3P 0.75 mM
G3P 1 mM
Si-Ctrl
Si-ENO1
Si-MTHFD1
Si-GSTP1
k

**Figure S7. Validation of the TRAP-assigned targets for an intrinsically reactive glycolytic metabolite glyceraldehyde 3-phosphate (G3P).**

**(a)** Administration of NaF (3 mM for 24 hr) and GNE140 (20 μM for 48 hr) suppressed the proliferation of HCT116 cells by CCK-8 assay. Data represent mean ± SEM (n = 5), two-tailed t-test, ***P < 0.001.

**(b)** Altered intracellular concentrations of glycolytic metabolites in response to NaF and GNE140 administration . Data represent mean ± SEM (n = 3), two-tailed t-test, *P<0.05,**P<0.01, ***P < 0.001, n.s. non-significant.

**(c)** HCT116 cell proliferation rate following administration of the accumulated glycolytic metabolites shown in (b). Data represent mean ± SEM (n = 5), two-tailed t-test, **P<0.01, ***P < 0.001.

**(d)** G3P significantly inhibited the proliferation of HT29 cells in a dose-dependent manner (n=5).

**(e-f)** Thermal shift assay shows MTHFD1 (e) and GSTP1 (f) of HCT116 cells both became stabilized by G3P incubation (30 μM for MTHFD1, 100 μM for GSTP1) in the tested temperature range compared to the vehicle.

**(g)** An iso-thermal dose-response experiment validated the G3P-induced stabilizing effect on ENO1 of HCT116 cells across the examined concentrations especially at 58.0 °C.

**(h)** TRAP ratios of the TRPs from ENO1 in HCT116 in response to G3P binding. Data represent mean ± SEM (n = 3), two-tailed t-test, **P<0.01, ***P < 0.001, ****P < 0.0001.

**(i)** Knockdown efficiency of siRNA used to selectively target ENO1, MTHFD1 and GSTP1 in HCT116 cells.

**(j)** G3P-induced proliferation suppression in control knockdown and ENO1/MTHFD1/GSTP1 knockdown HCT116 cells. Data represent mean ± SEM (n = 5), two-tailed t-test, *P < 0.05, **P < 0.01, ***P < 0.001, n.s. non-significant.

**(k)** Structural analysis of the TRAP-identified TRP shows G3P binding influenced the accessibility of the substrate-binding domain of GSTP1, since the minimal distance between the TRP (cyan, cartoon) and GSH (orange, sphere) is 7.69 Å (PDB: 5J41).

#### Figure 8

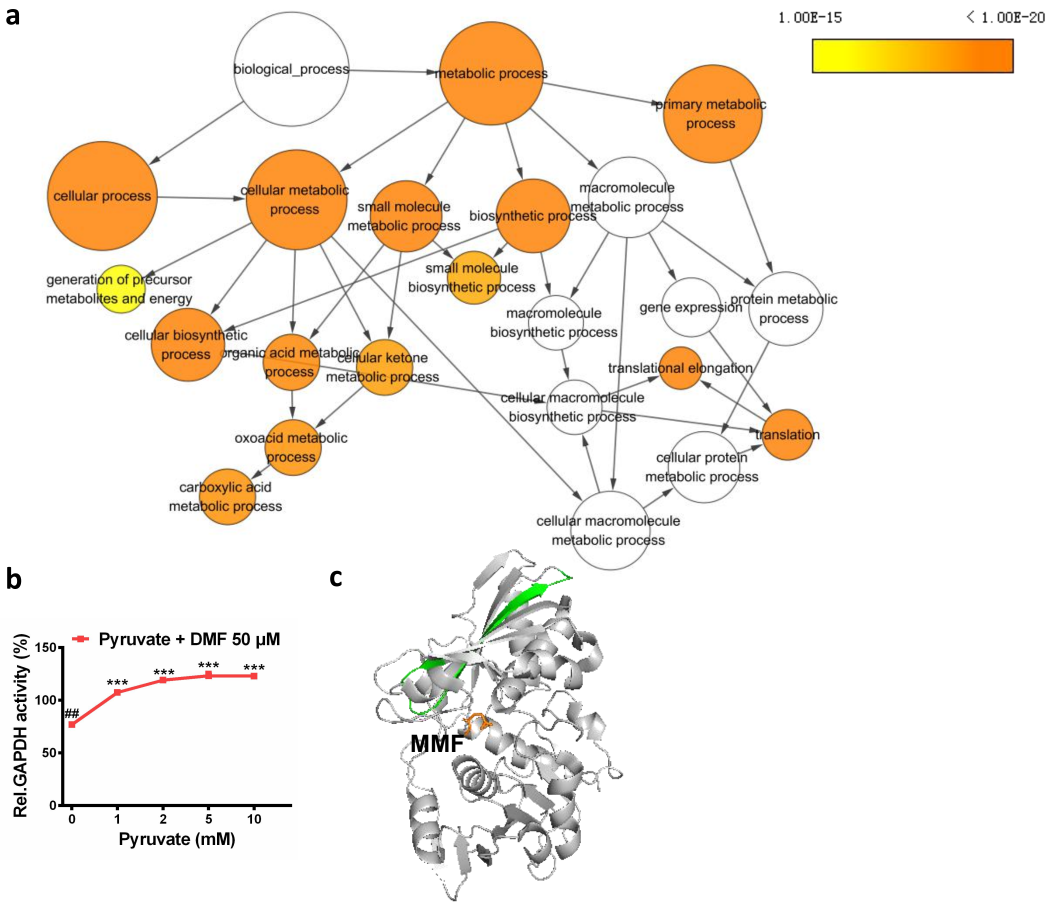


**Figure S8. Overall assessment of the druggability of the glycolytic metabolites targetome determined by TRAP from the HCT116 cells.**

**(a)** GO Biological Processes analysis of the druggable glycolytic targetome (that are classified into this category based on their availability in the DrugBank database).

**(b)** Administration of the newly identified GAPDH enzymatic activator, pyruvate, weakened the dimethyl fumarate (DMF)-induced GAPDH inhibition. Data represent mean ± SEM (n = 3), two-tailed t-test, $^{\#\#}$ P<0.01, ***P < 0.001. All comparisons were made between the GAPDH activity examined after the co-administration (pyruvate+DMF) and that with the DMF treatment alone.

**(c)** A K251-containing TRP (coded in green) in GAPDH (grey, cartoon) showing increased accessibility following pyruvate incubation as recorded in (Figure S4h) is located close to MMF (orange, stick) with a minimal Euclidean distance of 5.44 Å (PDB: 6IQ6). This suggests the loosened conformation of GAPDH spanning $V^{235}$-$K^{251}$ induced by pyruvate would weaken the engagement of MMF/DMF with GAPDH. DMF is supposed to bind to GAPDH in a similar fashion as MMF according to literature[22].

### Extended Data Supplementary Tables

**Table S1.** Summary of all glycolytic metabolites targetome discovered by TRAP from HCT116 cells.
**Table S2.** Summarized interactions of the known and quantified glycolytic metabolites-enzymatic targets as retrieved from BRENDA.
**Table S3.** Enriched KEGG pathway annotation network of the TRAP-identified glycolytic metabolites target proteins that are classified as those of catalytic activity..
**Table S4.** List of the minimal Euclidean distances measured between the detected TRPs from enzymatic targets that use these examined metabolites as substrates and their active sites.
